# Oncogene inactivation-induced senescence facilitates tumor relapse

**DOI:** 10.1101/2024.07.13.603369

**Authors:** Philipp Schmitt, Katrin Hönig, Maria Teresa Norcia, Marta F. Nogueira, Viktoria Flore, Inés Simó Vesperinas, Maria Villoro-Agud, Lushan Peng, Zuhal Safyürek, Mehreen Tariq, Ana Milojkovic, Kathleen Anders, Evelin Schröck, Sascha Sauer, Bora Uyar, Altuna Akalin, Gerald Willimsky, Simon Haas, Inmaculada Martínez-Reyes, Thomas Blankenstein

## Abstract

Oncogene-directed therapies can induce profound tumor regression in oncogene-addicted cancers such as BRAF-mutant melanoma and KRAS-driven pancreatic cancer, but their long-term benefit is often limited by resistance and early relapse. The mechanisms that allow residual cells to adapt, persist in a dormant state, and eventually fuel recurrence remain poorly understood. Here, we show that oncogene inactivation rapidly induces hallmark features of senescence together with a pro-inflammatory senescence-associated secretory phenotype (SASP). In vivo, oncogene inactivation-induced senescence (OIIS) predisposed tumors to relapse, characterized by polyploidy, chromosomal instability, and acquisition of alternative oncogenic pathways such as Mdm2 upregulation. Tumor microenvironment profiling by spectral flow cytometry revealed that relapse was associated with neovascularization and a shift from an immune-activated to an immunosuppressive milieu, indicating that senescent cells remodel their niche to promote regrowth. Importantly, OIIS features were also observed in human BRAF^V600E^ melanoma cells treated with vemurafenib, confirming the clinical relevance of our findings. Together, our findings establish OIIS as a double-edged process: it initially restrains tumor growth but simultaneously creates conditions that favor recurrence. By defining the genetic, metabolic, cytogenetic, and microenvironmental hallmarks of OIIS, our study highlights adaptations to oncogene deprivation that limit the durability of targeted therapies.

## INTRODUCTION

Oncogene-directed therapy has emerged as a promising approach in cancer drug development^1–4^. However, despite its potential, clinical efficacy is often limited by resistance and relapse^5–8^. Targeted therapies can induce profound tumor regression in oncogene-addicted cancers, but residual disease often persists, eventually giving rise to recurrence. The mechanisms that enable cancer cells to adapt to oncogene inactivation, survive in a dormant state, and resume proliferation remain incompletely understood. In this study, we investigated these processes using a mouse cancer model in which cancer cell proliferation is driven by the oncogene SV40 large T antigen (Tag) in an inducible manner, allowing us to activate and inactivate the oncogene *in vitro* and *in vivo*. Importantly, this system mimics clinical scenarios where tumor growth is dependent on oncogene signaling and recapitulates oncogene addiction, a phenomenon well-documented in human cancers such as BRAF-mutant melanoma and KRAS-driven pancreatic cancer. Our results show that oncogene inactivation leads to the acquisition of cellular senescence and induction of a pro-inflammatory senescence-associated secretory phenotype (SASP). Senescence is widely defined as a durable form of growth arrest and is linked to extensive cellular stress that can be induced by different types of insults, including oncogenic activation, radiation, or chemotherapy^9^. Since the seminal work of Hayflick and Moorhead in 1961, our understanding of senescence has expanded, with increasing evidence for both beneficial and detrimental roles in aging and disease^10–12^.

In the cancer context, senescence plays a complex, dual role. Oncogene-induced senescence (OIS) is an early barrier to malignant transformation, enforcing cell-cycle arrest and, in some models, initiating immune-mediated clearance through SASP-driven recruitment of effector cells^13–15^. However, if senescent cells persist, their SASP can paradoxically remodel the tumor microenvironment, promoting inflammation, extracellular matrix degradation, and the secretion of growth-promoting factors that facilitate tumor progression^16–18^. Therapy-induced senescence (TIS) has historically been viewed as a favorable outcome of cancer treatment, suppressing tumor growth by arresting the proliferation of genetically unstable cancer cells^19^. However, mounting evidence suggests that TIS may instead promote long-term tumor progression. Cancer cells that survive therapy often acquire senescent features and may later escape this state, leading to more aggressive relapses^20–22^. In acute myeloid leukemia, chemotherapy induces a transient senescence-like state from which cells re-enter the cell cycle with enhanced stemness and resilience^22^. Similarly, in B-cell lymphoma, TIS leads to epigenetic reprogramming toward stemness, increasing tumor-initiating capacity^21^. Over time, senescent tumor cells can accumulate epigenetic “scars” that predispose them to senescence escape, chromatin remodeling, and oncogenic reactivation^23^. Beyond classical inducers of senescence, a growing body of work indicates that targeted oncogene inactivation itself can trigger senescence in oncogene-addicted tumors. This phenomenon has been described following c-MYC inactivation in lymphoma^24^, after BRAF inhibition in melanoma^25^ and after KRAS inhibition in pancreatic cancer^26^. Whether OIIS enforces durable growth arrest or instead creates a state prone to relapse has remained unclear.

Here, we address this gap using a controllable oncogene switch model in which SV40 large T antigen (Tag) drives tumor cell proliferation in an inducible manner. This system recapitulates oncogene addiction and allows us to study the effects of oncogene activation and inactivation *in vitro* and *in vivo*. Using epithelial (clone 4 gastric carcinoma) and mesenchymal (TTC#3055 spindle cell sarcoma) tumor cell lines, we demonstrate that OIIS predisposes tumors to relapse and is associated with adverse outcomes. We further explore how OIIS shapes tumor evolution, including chromosomal instability, acquisition of alternative oncogenic pathways, and metabolic adaptations. Finally, we validate key features of OIIS in human BRAF^V600E^ melanoma cells, confirming that our findings are relevant to oncogene-driven cancers.

## RESULTS

### Oncogene inactivation induces senescence and SASP

To study the mechanisms enabling cancer cells to adapt to oncogene inactivation, we used cancer cells isolated from tumors that grew sporadically in transgenic mice ^27,28^. Specifically, we used cells from a gastric carcinoma (clone 4) and from a spindle cell sarcoma isolated from the snout (TTC#3055). These established tumor cell lines express the oncogene Tag fused to the reporter gene firefly luciferase (TagLuc) under the tetracycline response element (TRE) together with the rtTA2S-M2 transactivator (rtTA). In clone 4 cells, the loxP-flanked stop cassette was excised by transient gene transfer in cultured cells with a Cre recombinase-encoding adenovirus. This system allows conditional activation or inactivation of the oncogene by adding or withdrawing doxycycline (dox), respectively, both *in vitro* and *in vivo*. Fusion of Tag to luciferase enables monitoring of tumor cell growth *in vivo* by bioluminescence imaging upon subcutaneous transplantation (Figure 1A). This model provides an ideal setting to analyze the effects of oncogene inactivation on cancer cells and tumor development. We first confirmed loss of TagLuc expression, indicating oncogene inactivation, as early as one day after dox withdrawal in clone 4 and in TTC#3055 cells (Figure 1B and Extended Data Figure 1A). Following oncogene inactivation, cellular proliferation was halted, and cells progressively exhibited cell cycle arrest (Figure 1C, D, Extended Figure 1B). Cells became enlarged, flattened, and displayed an irregular morphology, characteristic of a senescent phenotype (Figure 1E and Extended Data Figure 1C). Following TagLuc inactivation, a significant increase in senescence-associated β-galactosidase (SA-β-gal) activity was detected in clone 4 cells (Figure 1E and Extended Data Figure 1C). To further validate senescence induction, we analyzed the expression of Cdkn1a (p21) and Cdkn2a (p16), two key cell cycle regulators associated with senescence^29^. Consistent with senescence acquisition, we observed increased p21 protein levels after oncogene inactivation (Extended Data Figure 1E). However, p16 protein was downregulated, a finding that deviates from the canonical senescence signature (Extended Data Figure 1E). Analysis of mRNA levels confirmed a significant upregulation of p21 and a downregulation of p16 following TagLuc inactivation (Extended Data Figure 1F). A hallmark of senescent cells is the senescence-associated secretory phenotype (SASP), characterized by cytokine and matrix-remodeling enzyme secretion^17^. We analyzed the SASP by generating conditioned medium from proliferating (TagLuc-expressing) and senescent (TagLuc-negative) clone 4 cells. Using a semi-quantitative cytokine array, we observed an increase in the secretion of interleukin-6 (Il-6), matrix metalloprotease 2 (Mmp2), and matrix metalloprotease 3 (Mmp3) in senescent cells (Extended Data Figure 1G). ELISAs confirmed elevated Il-6 and Mmp3 secretion after senescence induction (Figure 1F, G).

**Figure 1:**
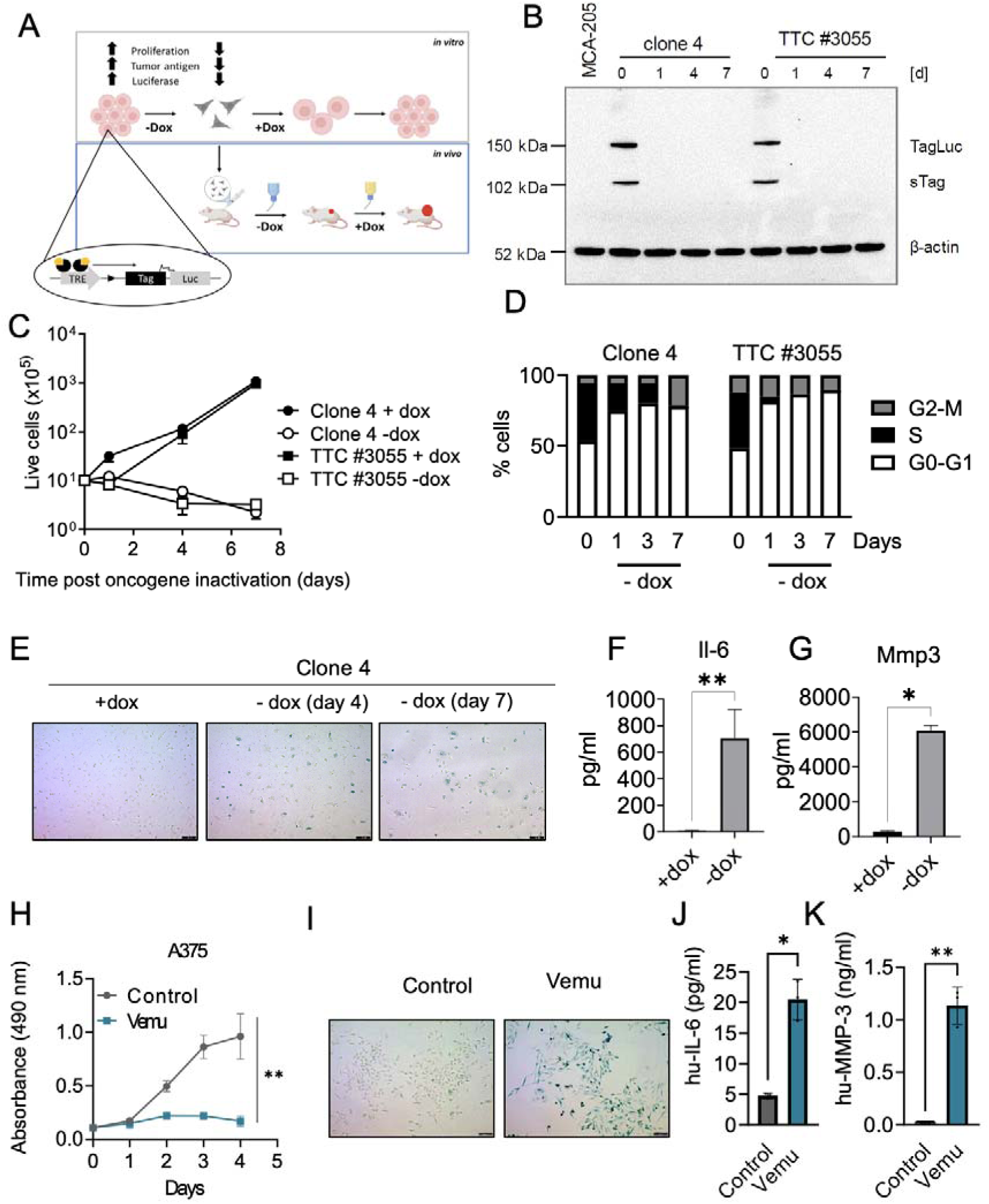
Oncogene inactivation induces senescence and a pro-inflammatory SASP. **A,** Schematic of the mouse model. The model features the SV40 large T antigen (Tag) oncogene regulated by a tetracycline response element (TRE). Addition or withdrawal of doxycycline (dox) activates or inactivates the oncogene, respectively. The Tag is fused to firefly luciferase (Luc), creating the TagLuc fusion protein, which serves as both a cancer driver and a reporter for intravital imaging. Oncogene activation and inactivation are reversible both *in vitro* and *in vivo*. **B,** Western blot analysis of TagLuc in clone 4 and TTC#3055 cells cultured in the presence of dox or at different time points after dox withdrawal. β-actin was used as a loading control. MCA-205 served as negative control. A shorter splicing product of Tag (∼ 100 kDa), sTagLuc was detected. Shown is one representative experiment out of four independent experiments. **C,** Clone 4 and TTC#3055 cells were grown in the presence or absence of dox and live cell number was assessed at different time points. Error bars indicate standard deviation; n=2 technical replicates. **D,** Cell-cycle phase distribution was determined by BrdU incorporation following TagLuc inactivation. Data shown are from one representative experiment out of three independent experiments. **E,** SA-β-galactosidase staining of clone 4 cells grown on dox or after four or seven days post-TagLuc inactivation. Scale bar, 100 μm. **F, G** Clone 4 cells were cultured with or without dox for two weeks. Conditioned medium was generated and normalized to cell number (2×10^5^ cells/ml). Il-6 (F) and Mmp3 (G) secretion was analyzed via ELISA. Error bars indicate standard deviation. Il-6: n = 4, pooled data from two independent experiments comprising two samples, ** p < 0.01. Mmp3: n = 2 technical replicates from one experiment, * p < 0.05. **H,** Proliferation of A375 melanoma cells treated with vemurafenib (Vemu, 1 µM) or vehicle control was measured by MTS assay at the indicated time points (mean ± s.d.; n = 3). **p < 0.01. **I,** Bright-field microscopy of A375 cells treated with Vemu showing flattened morphology, enlarged cell shape and positive SA-β-galactosidase staining. Scale bar, 100 μm **J, K** Secreted SASP factors hu-IL-6 (J) and hu-MMP-3 (K) measured by ELISA in the supernatant of control vs. Vemu-treated cells for 15 days (mean ± s.d.; n = 3).

To confirm the relevance of our observations in a human context, we next assessed whether therapeutic oncogene inhibition induces senescence and SASP in human melanoma cells. Treatment of A375 cells with the mutant BRAF inhibitor vemurafenib led to a strong reduction in proliferation compared to vehicle-treated controls (Figure 1H). Vemurafenib-treated cells acquired classical features of senescence, including SA-β-gal staining and a flattened, enlarged morphology (Figure 1I). Immunoblotting revealed a progressive decrease in phosphorylated Rb (pRb) without upregulation of p21 or p16 (Extended Figure 1I, J), consistent with the acquisition of a non-canonical senescence program reported in melanoma cells upon mutant BRAF inhibitors treatment^37^. To evaluate SASP induction in human cells, we measured the secretion of IL-6 and MMP-3 in A375 cells treated with vemurafenib. Both factors were upregulated upon treatment, confirming the acquisition of a SASP in this model (Figure 1J, K).

Collectively, the concordant findings in mouse and human cells show that oncogene inactivation leads to the acquisition of a senescent phenotype.

### TagLuc inactivation-mediated senescence promotes major transcriptomic changes and the acquisition of a plurimetabolic state

To further characterize the phenotype of OIIS, we performed mRNA sequencing of clone 4 cells in two conditions: proliferating (TagLuc-expressing, +dox) and senescent (TagLuc-negative, -dox). Differential gene expression analysis revealed profound transcriptional reprogramming, with 1,067 genes significantly upregulated (p-adj < 0.001, log2 fold-change > 2) and 928 genes significantly downregulated (p-adj < 0.001, log2-fold-change < -2) in senescent cells compared to proliferating cells (Extended Figure 2A; Supplementary Document 1). Gene set enrichment analysis (GSEA) demonstrated that pathways related to cell cycle regulation, mitosis, and DNA replication were among the most significantly downregulated in senescent cells, consistent with the G0/G1 arrest observed after oncogene inactivation (Figure 2A). Conversely, senescent cells showed marked upregulation of pathways involved in extracellular matrix (ECM) remodeling, including keratan sulfate degradation, ECM-receptor interaction, collagen degradation, and matrix organization (Figure 2B). Additionally, senescent cells exhibited enrichment of inflammatory signaling and growth factor-related pathways such as cytokine-cytokine receptor interactions, complement activation, PI3K-AKT signaling, and insulin-like growth factor (IGF) transport (Figure 2B). Sphingolipid metabolism, which has been previously implicated in the regulation of senescence, was also upregulated, reflecting changes in membrane dynamics and metabolic adaptation^30^. The senescence-associated secretory phenotype (SASP) was confirmed by increased expression of *Mmp13*, *Mmp3*, *Igfbp2*, *Il6*, and other inflammatory mediators (Extended Figure 2B).

**Figure 2:**
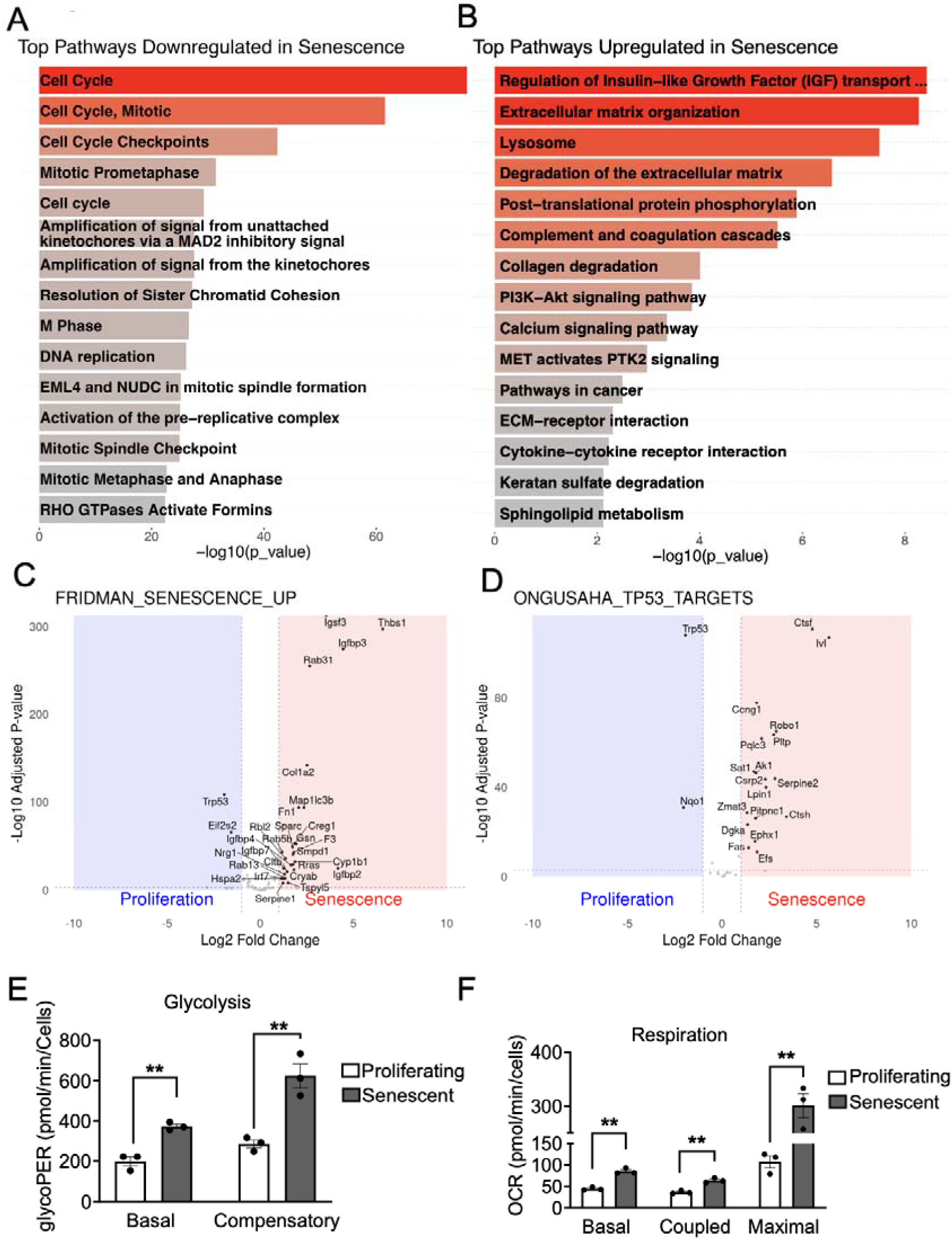
Senescence is accompanied by major transcriptional changes and the acquisition of a plurimetabolic state. **A, B,** Enriched pathway terms (KEGG/Reactome) among the downregulated (A) and upregulated (B) genes in senescent (TagLuc-negative, -dox) cells compared to proliferating (TagLuc-expressing, +dox) cells. The complete dataset of differentially expressed genes can be found in Supplementary Document 1. n=3 technical replicates. **C, D,** Gene-set enrichment analysis using the FRIDMAN_SENESCENCE_UP and ONGUASAHA_TP53_TARGETS gene sets in senescent (TagLuc-negative, –dox) versus proliferating (TagLuc-expressing, +dox) clone 4 cells. n = 3 technical replicates. **E,** Basal and compensatory glycolysis in senescent (TagLuc-negative, -dox) cells compared to proliferating (TagLuc expressing, +dox) cells. Data are mean ± s.e.m; n = 3 independent experiments. *p < 0.05, two-tailed t-tests. **F,** Basal, coupled, and maximal respiration in senescent (TagLuc-negative, -dox) cells compared to proliferating (TagLuc-expressing, +dox) cells. Data are mean ± s.e.m; n = 3 independent experiments. *p < 0.05, two-tailed t-tests.

Consistent with senescence induction, the FRIDMAN_SENESCENCE_UP gene set was significantly enriched in senescent cells (Figure 2C)^31^. We next analyzed p53 pathway activity using the ONGUASAHA_TP53_TARGETS gene set, which revealed selective enrichment of senescence-associated p53 targets in senescent cells (Figure 2D)^32^. Analysis of the Cancer Single-Cell State Atlas (CancerSEA) revealed that senescent cells downregulate cell cycle, DNA repair, DNA damage response, and proliferation signatures, while upregulating pathways linked to hypoxia, angiogenesis, invasion, and inflammation (Extended Figure 2C, D)^33^.

In addition, we analyzed the metabolic phenotype of proliferating versus senescent clone 4 cells. Senescent cells displayed a higher basal glycolytic rate, as well as increased compensatory glycolysis after mitochondrial inhibition with rotenone and antimycin A (Figure 2E; Extended Figure 2E). We also observed that basal, coupled, and maximal oxygen consumption rates were significantly higher in senescent cells compared to proliferating controls, indicating enhanced mitochondrial respiration (Figure 2F; Extended Figure 2F). To functionally test the dependency of senescent cells on glycolysis and respiration, we treated cells with 2-deoxyglucose (2-DG) or rotenone/antimycin A (R/A) individually. Senescent cells displayed increased sensitivity to both glycolysis inhibition and mitochondrial respiration blockade compared to proliferating cells, with a significant increase in cell death upon either treatment (Extended Figure 2G). This finding supports the dual activation of glycolysis and oxidative phosphorylation, confirming that senescent cells acquire a plurimetabolic state for survival.

### Oncogene inactivation-induced senescence predisposes to cancer relapse

To analyze how OIIS influences tumor relapse, we established three complementary *in vivo* conditions (Figure 3): a) a never-senescent setting with continuous oncogene expression (+dox throughout); b) a transient OIIS setting in which TagLuc is switched off and later reactivated (dox off → on), and c) a long-term OIIS setting in which TagLuc remains suppressed for the entire experiment (dox off throughout). These three conditions allow us to compare tumors that never experience senescence, tumors that undergo a reversible senescent phase, and tumors under a long-term senescent state.

**Figure 3:**
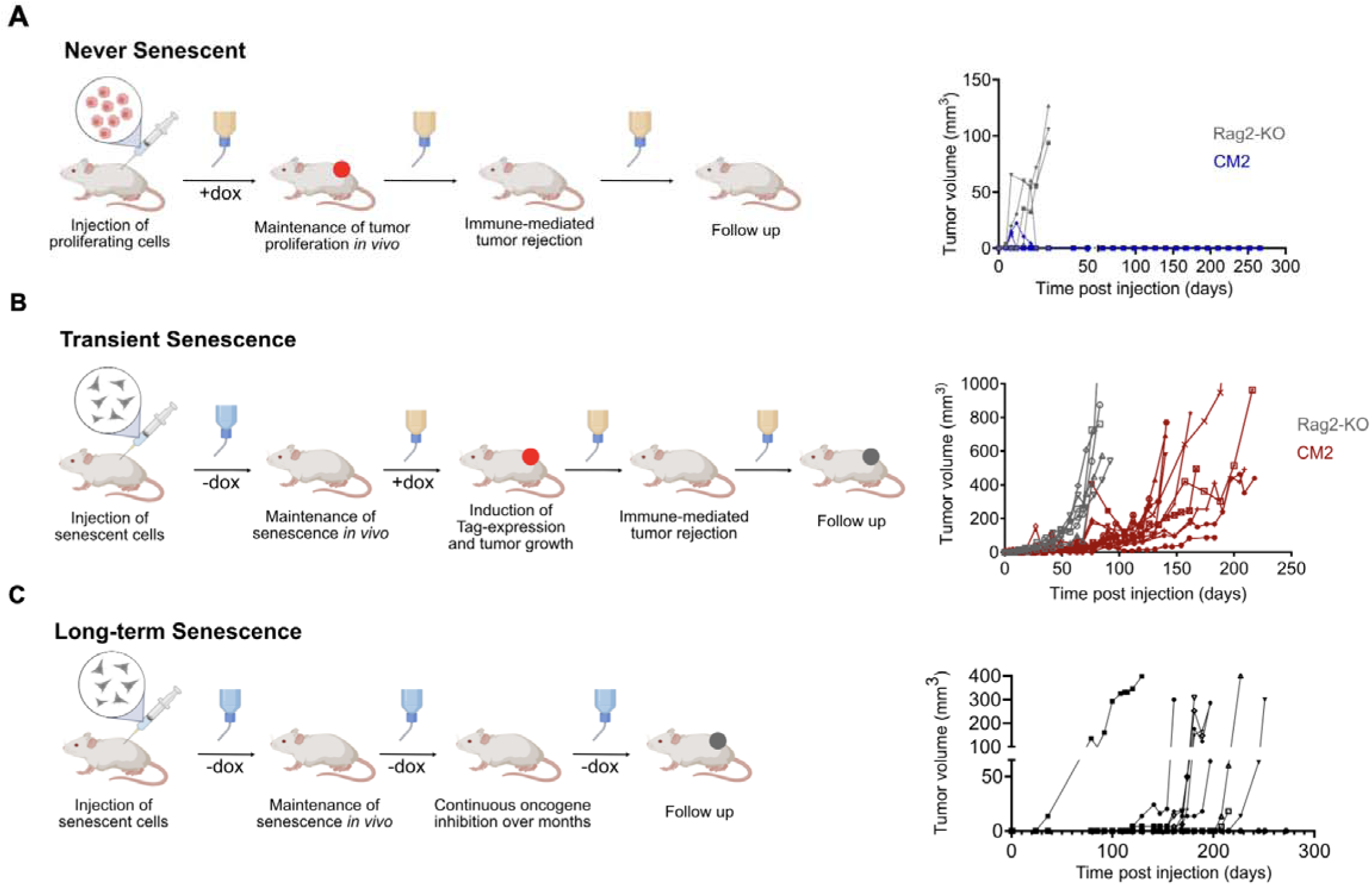
Distinct senescence regimens shape tumor regression and relapse in vivo. **A,** *Never-senescent condition*. Schematic (left) and tumor growth curves (right) for mice injected s.c. with proliferating clone 4 cells (TagLuc-expressing, +dox) and maintained on doxycycline throughout. Continuous +dox preserves TagLuc expression and tumor cel proliferation in both CM2 and Rag2-KO mice. In immunocompetent CM2 mice (blue), tumors initially grow but are subsequently eliminated by Tag/SV40-specific immune responses, whereas in Rag2-KO mice (grey) tumors grow progressively. No tumor relapse is observed in CM2 mice during long-term follow-up. **B,** *Transient OIIS condition.* Schematic (left) and tumor growth curves (right) for mice injected with senescent clone 4 cells generated by 14 days of dox withdrawal *in vitro* (dox off). After injection, mice are kept off dox for four weeks to maintain OIIS *in vivo*, and then switched to +dox to reactivate TagLuc and induce tumor growth. In CM2 mice (red), tumors initially expand under +dox and are then rejected by the immune system, but after a prolonged latency tumors relapse. In Rag2-KO mice (grey), tumors grow continuously without regression, consistent with the absence of adaptive immunity. **C,** *Long-term OIIS condition*. Schematic (left) and tumor growth curves (right) for mice injected with senescent clone 4 cells and kept off dox for the entire duration of the experiment, resulting in sustained oncogene inactivation *in vivo*. Despite continuous TagLuc suppression, tumors eventually arise in most CM2 mice (black) after a long latency, indicating senescence escape and oncogene-independent outgrowth.

First, we injected proliferating clone 4 cells (TagLuc expressing, +dox) into CM2 and Rag2-KO mice, while maintaining dox throughout (Figure 3A). CM2 mice ubiquitously express the rtTA transactivator and are therefore tolerant to it, eliminating the possibility of immune responses against the transactivator itself. Tumor burden was quantified by tumor volume and TagLuc-derived bioluminescent flux, the latter directly reporting the *in vivo* burden of oncogene-expressing tumor cells, as the firefly luciferase is fused to the oncogene Tag. In CM2 mice, tumors initially grew but were consistently rejected by immune responses targeting TagLuc shown by staining of peptide IV-MHC I tetramer^+^ CD8⁺ T-cells (Figure 3A and Extended Figure 3A, B). Importantly, no tumor relapse was observed in these mice over a long observation period. In Rag2-KO mice, tumors grew progressively consistent with the absence of immune-mediated clearance (Figure 3A, Extended Figure 3E).

Next, we withdrew dox from cultured clone 4 cells for 14 days to induce senescence *in vitro* (Figure 3B). Senescent cells were then injected into immunocompetent CM2 mice and immunodeficient Rag2-KO mice. After injection, the cells were kept in a senescent state *in vivo* by maintaining mice off dox for four weeks. We then introduced dox via the drinking water to reactivate TagLuc expression and stimulate tumor growth. This design therefore models a transient phase of OIIS, in which senescence is imposed during dox withdrawal (*in vitro* and *in vivo*) and is reversed upon reactivation of TagLuc once dox is introduced *in vivo*. In CM2 mice, tumors initially grew after dox reintroduction but subsequently regressed due to immune-mediated rejection, as confirmed by bioluminescence imaging and measurement of tumor volume (Figure 3B and Extended Figure 3C). Re-introduction of dox restored TagLuc expression, rendering tumor cells immunogenic and leading to their elimination by Tag/SV40-specific CD8⁺ T cells (Extended Figure 3D). In Rag2-KO mice, by contrast, tumors grew progressively without regression (Figure 3B, Extended Figure 3F). Notably, after a prolonged latency, tumor relapse occurred in all CM2 mice injected with transiently senescent clone 4 cells. Similar results were obtained with TTC#3055 cells: after oncogene reactivation *in vivo*, tumors in CM2 mice initially grew, regressed, and later relapsed, whereas in Rag2-KO mice, tumors grew continuously (Extended Figure 3G, H).

To summarize, continuous oncogene expression in the absence of any senescent phase leads to durable immune-mediated eradication (in CM2 mice) but not to relapse. A preceding transient phase of OIIS, on the other hand, leads to tumor relapse after initial regression. These findings indicate that it is specifically a preceding transient OIIS phase that creates a unique condition which predisposes tumors to late relapse despite initial immune clearance.

We next tested whether long-term senescence alone could lead to tumor formation without oncogene reactivation (Figure 3C). In contrast to the transient OIIS protocol above, where dox withdrawal and re-addition impose a reversible senescent phase, here oncogene expression remains suppressed throughout the entire experiment. Clone 4 cells maintained off dox for 14 days were injected into CM2 mice, which were then kept continuously off dox. Remarkably, tumors developed in 10 out of 12 mice after a long latency, despite continuous oncogene suppression. This indicates that a subset of OIIS cells can eventually undergo “senescence escape”, i.e. lose their growth arrest and re-enter the cell cycle *in vivo*, even without reactivation of the original oncogene. To explore the underlying mechanism, we re-isolated cell lines from relapsed tumors in both experimental conditions (transient and long-term senescence). In four of five cell lines, TagLuc expression was undetectable by luciferase activity and mRNA analysis (Extended Figure 3I; J), consistent with the absence of bioluminescence in several relapsed tumors. Thus, tumor outgrowth after OIIS can occur independently of the original oncogene, suggesting selection for alternative oncogenic pathways. Together, these results demonstrate that OIIS creates a cellular state that fosters oncogene bypass and tumor relapse.

To determine whether vemurafenib-induced OIIS is transient, we first treated A375 melanoma cells with vemurafenib in vitro to induce growth arrest and acquisition of senescence markers. Upon withdrawal of vemurafenib, cells resumed proliferation (Extended Figure 3K), indicating that the senescent state induced by mutant BRAF inhibition is reversible. In a complementary in vivo experiment, we pretreated A375 cells with vemurafenib *in vitro* to induce senescence, implanted them subcutaneously, and continued vemurafenib treatment of recipient mice for four weeks (Extended Figure 3L). As shown in Extended Figure 3M, tumors formed and grew despite this extended senescence-inducing regimen, further supporting the transient nature of therapeutic OIIS in this model. These findings demonstrate that BRAF inhibitor–induced OIIS in human melanoma cells is not durably tumor-suppressive but instead creates a state that permits senescence escape and tumor regrowth. Together, these results show that OIIS creates a cellular state that fosters oncogene bypass and tumor relapse.

### Cancer cells from relapsed tumors depend on Mdm2 and display polyploidy

To investigate the molecular mechanisms supporting tumor relapse after OIIS, we analyzed transcriptional and cytogenetic features of cancer cell lines derived from relapsed tumors. We compared these lines to the parental proliferating clone 4 cells cultured in the presence of dox and to senescent clone 4 cells maintained off dox. Transcriptomic profiling revealed 143 genes significantly upregulated and 67 genes downregulated in relapsed cells relative to both parental and senescent populations (Supplementary Document 2). Pathway enrichment analysis of these differentially expressed genes indicated that relapsed tumors acquired molecular signatures associated with enhanced proliferation and growth factor signaling. Among the top upregulated pathways were MAPK signaling, RAF/MAPK kinase cascades, PI3K-AKT signaling and cytokine-receptor interactions (Figure 4A). This profile reflects the activation of pro-growth signaling networks that are typically suppressed during senescence. Conversely, genes related to extracellular matrix organization, collagen-containing structures, and cellular adhesion were significantly downregulated in relapsed tumors (Figure 4B). This shift suggests that relapsed tumors not only reactivate proliferative programs but also remodel their microenvironment, potentially facilitating tumor growth. Of note, gene set enrichment analysis comparing relapsed samples to the senescent state showed that relapsed tumors still show enrichment of senescence-related genes among the genes upregulated (Extended Figure 4A). These findings suggest that relapsed cells acquire a hybrid state, maintaining residual senescence features, while engaging proliferation and growth signaling programs. We next assessed the metabolic changes in relapsed tumors, which showed a marked shift in metabolic pathway usage (Extended Figure 4B). While glycosaminoglycan metabolism, sphingolipid metabolism, and keratan sulfate metabolism were downregulated, relapsed cells upregulated nucleotide metabolism, amino acid metabolism, and folate cycle pathways, reflecting increased anabolic activity to support proliferation (Extended Figure 4B).

**Figure 4:**
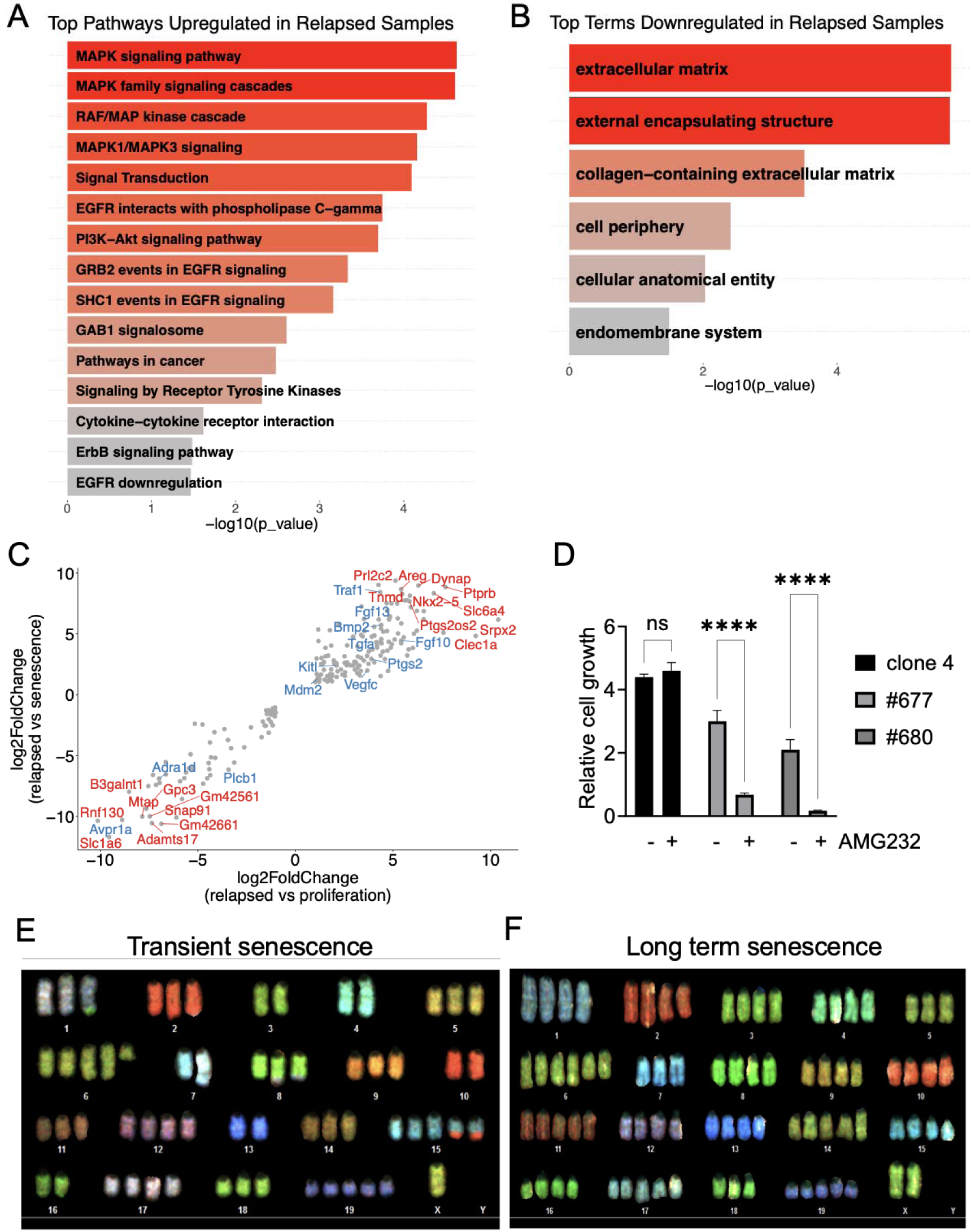
Cancer cells re-isolated from relapsed tumors depend on Mdm2 and acquired polyploidy. **A, B,** Top enriched pathways among genes upregulated (A) or downregulated (B) in relapse-derived cancer cell lines compared to both parental proliferating (TagLuc-expressing, +dox) and senescent (TagLuc-negative, –dox) clone 4 cells. Pathway enrichment was based on significantly differentially expressed genes (adjusted p < 0.05, |log₂FC| > 1). **C,** Genes that are upregulated and downregulated in relapsed cancer cell lines compared to proliferating (TagLuc-expressing, +dox) and senescent (TagLuc-negative, -dox) clone 4 cells. Genes in red represent the top 10 most strongly up- or downregulated genes across both comparisons; genes in blue are additional relevant candidates preselected as potential therapeutic targets. **D,** Relative cell growth of parental clone 4 cells and cancer cell lines from relapsed tumors (#677, #680) following treatment with the Mdm2 inhibitor AMG-232 (1 µM for 72 hours). Cell growth was calculated by comparing the cell number after treatment to the initial cell number. Data are shown as mean ± SD (n = 3), \*\*\*\**p* < 0.0001. **E, F,** Representative spectral karyotyping (SKY) images of relapse-derived cell lines from tumors that emerged after transient TagLuc inactivation (E) or long-term TagLuc inactivation (F). Images were obtained from 8–10 randomly selected metaphase spreads per cell line, with one representative example shown. Supplementary Document 4 details the chromosomal aberrations observed in the 10 selected metaphase chromosomes of the parental Clone 4 cells.

Among the upregulated genes in relapsed tumors, Mdm2 emerged as a potential oncogenic driver (Figure 4C). Since Mdm2 is a key negative regulator of Tp53, its increased expression could compensate for the loss of Tag-mediated Tp53 suppression following oncogene inactivation. To test this hypothesis, we treated relapsed tumor-derived cell lines with the Mdm2 inhibitor AMG-232^34^. The growth of relapsed cells was significantly impaired upon Mdm2 inhibition, whereas the parental clone 4 cells remained unaffected (Figure 4D). These results confirm that relapsed tumors acquire an Mdm2-dependent proliferative advantage.

To evaluate whether genomic instability contributes to relapse, we performed multicolor karyotyping on the tumor-derived cell lines. Parental clone 4 cells maintained under continuous oncogene expression exhibited a near-diploid karyotype (Extended Figure 4C). In contrast, and in agreement with previous findings, relapsed tumor-derived cells displayed extensive chromosomal abnormalities and polyploidy (Figure 4E, F and Extended Figure 4D-F)^35,36^. Both transient- and long-term-senescence–derived relapsed cells exhibited dramatic chromosomal rearrangements and increased chromosome numbers, consistent with the acquisition of highly polyploid genomes. Despite the restoration of Rb and p53 function after Tag inactivation, these relapsed tumors retained complex structural aberrations and genomic instability. Altogether, these findings reveal that tumor relapse after OIIS is associated with a combination of genomic instability, metabolic reprogramming, and the activation of alternative oncogenic pathways such as Mdm2 upregulation.

### Distinct immune microenvironment landscapes characterize progressive tumors, remission and relapse

To investigate how the tumor microenvironment contributes to OIIS during remission and to the subsequent relapse, we analyzed the immune and stromal cell populations across three experimental groups of Rag2-KO tumor-bearing mice. In the first group, tumors were maintained in a proliferative state through continuous dox administration, allowing persistent oncogene expression and progressive tumor growth. In the second group, once tumors were established, dox was withdrawn to inactivate the oncogene and induce remission. In the third group, following a remission phase induced by dox withdrawal, oncogene expression was reactivated to mimic relapse, providing a model to study the microenvironment during tumor regrowth. A schematic of this experimental setup is shown in Figure 5A. Longitudinal bioluminescence imaging and tumor measurements confirmed progressive tumor growth upon sustained oncogene expression; tumors in the remission group regressed following oncogene inactivation; and tumors in the relapse group re-emerged after oncogene reactivation (Figure 5B–C and Extended Figure 5A).

**Figure 5:**
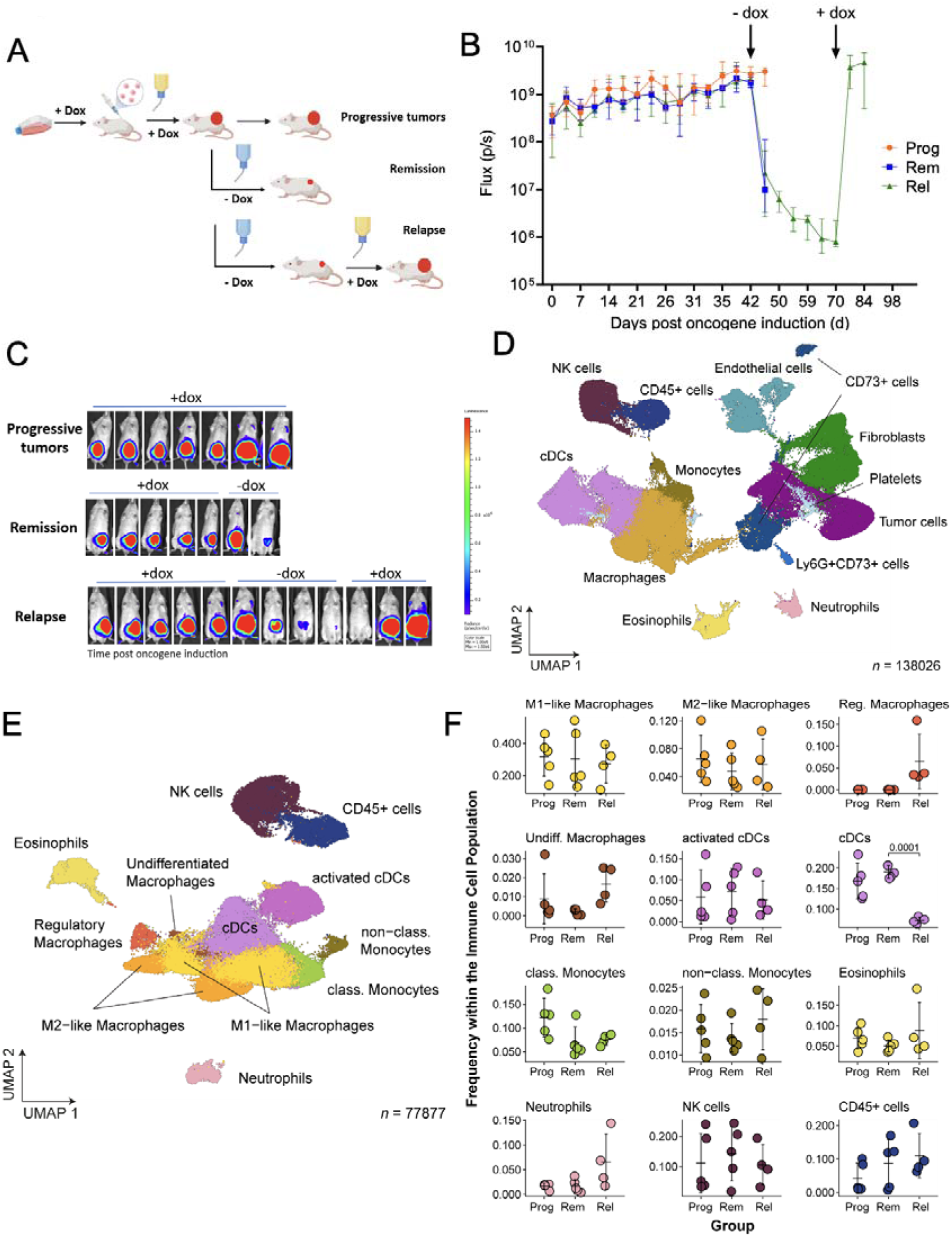
Immune infiltration in progressive tumors, remission and relapse. **A,** Schematic representation of the OIIS mouse cancer model and relapse design. Clone 4 cancer cells cultured in the presence of doxycycline (dox) were subcutaneously injected (1 × 10l cells in Matrigel) into Rag2-KO mice. Dox (200 µg/ml) was provided in the drinking water at tumor inoculation. In the remission and relapse groups, dox was withdrawn once tumors reached a clinically relevant size to induce OIIS. In the relapse group, oncogene expression was re-induced by re-administering dox once the BL signal was no longer detectable. **B,** BL kinetics of Rag2-KO mice in the progressive (Prog; n=5), remission (Rem; n=5) and relapse (Rel; n=5) groups. Dox was removed at day 42 from the remission and relapse groups and re-administered at day 70 in the relapse group. Tumor plugs were isolated at day 45 from the progressive and remission groups and at day 102 from the relapse group. **C,** Representative BL images of Rag2-KO mice from each group (exposure time: 60 s). **D,** UMAP representation of the overall cellular landscape. Recorded cells were processed with PICtR; out of 1,024,994 cells, 138,026 sketched cells are displayed. **E,** UMAP representation of immune cells. Out of 318,115 cells, 77,877 sketched cells are displayed. **F,** Frequency of the immune cell types identified in panel E within each experimental group. *P* values were determined with a two-sided Welch’s *t*-test and corrected according to Benjamini-Hochberg. Error bars indicate the mean and standard deviation. cDCs: conventional dendritic cells, NK cells: natural killer cells, non-class./class. Monocytes: non-classical/classical monocytes, UMAP: Uniform Manifold Approximation and Projection.

To characterize the tumor microenvironment across progression, remission and relapse, we performed spectral flow cytometry of extracted tumor plugs (tumor cells were always injected in matrigel). Uniform Manifold Approximation and Projection (UMAP) visualization revealed the global composition of the tumor microenvironment, including immune, stromal, and endothelial populations that were identified using lineage markers (Figure 5D and Extended Figure 5B). UMAPs stratified by condition highlighted shifts in the cellular composition of relapsed tumors (Extended Figure 5C). Interestingly, we observed a trend towards reduced endothelial cells upon induction of remission, followed by a significant increase in relapsed tumors (Extended Figure 5D), likely reflecting transient vascular disruption after extensive tumor cell death following oncogene inactivation and subsequent neovascularization during tumor regrowth. Focusing on the immune compartment, we identified diverse cell types including NK cells, conventional and activated dendritic cells (cDCs), M1-like and M2-like macrophages, regulatory and undifferentiated macrophages, eosinophils, neutrophils and classical and non-classical monocytes (Figure 5E and Extended Figure 5E). Relapsed tumors displayed the most notable changes in the relative abundance of these populations (Figure 5F and Extended Figure 5F). In particular, we observed a significant reduction in cDCs and an increase in regulatory macrophages in relapsed tumors, pointing to a shift towards a more immunosuppressive microenvironment (Figure 5F). These findings indicate that tumor relapse after oncogene inactivation is not solely driven by cell-intrinsic resistance mechanisms but also shaped by dynamic remodeling of the immune microenvironment.

## DISCUSSION

In this study, we analyzed the effects of oncogene inactivation *in vitro* and *in vivo* and found that acquisition of cellular senescence, together with a pro-inflammatory SASP and genetic and metabolic alterations, are key adaptive mechanisms. Our results align with a broader spectrum of work showing that suppression of frequent oncogenes such as c-MYC, BRAF, or KRAS induces senescence programs that contribute to tumor regression but can also promote therapy resistance^25,41–44^. In our models, oncogene inactivation induced hallmark features of senescence, including cell cycle arrest, enlarged morphology, and SA-β-gal activity. Yet, unlike canonical senescence, p16 was consistently downregulated. In our mouse model system, this is explained by the regulatory interplay between Tag, p53, and Rb. SV40 Tag binds and functionally inactivates both p53 and Rb^45–47^, thereby preventing stress-induced apoptosis and cell-cycle arrest^48^. Tag binding stabilizes p53 protein but abolishes its transcriptional activity. Upon Tag withdrawal, this inhibition is released, resulting in selective enrichment of p53 target gene expression despite reduced p53 levels, creating a p21-dependent, but p16-independent senescence state. The downregulation of p16 is consistent with a feedback loop whereby Rb negatively regulates p16 transcription. In Rb-deficient or Tag-expressing cells, p16 accumulates due to loss of repression; conversely, restoration of Rb function when Tag is switched off suppresses p16 expression^49,50^. Thus, our system reflects how viral oncogene withdrawal reshapes tumor suppressor regulation, shifting senescence away from canonical p53–p16 signaling. We found that a similar noncanonical senescence program operates in human melanoma cells. In A375 BRAF^V600E^ melanoma cells, p16 also failed to increase upon oncogene inactivation, consistent with previous findings and the frequent p16 loss in melanoma cell lines^37,51^. Instead, vemurafenib-induced senescence was mediated by cyclin D1/p-Rb inhibition without robust activation of the DNA damage/p53/p21 axis in A375 cells. Together, these findings highlight that senescence can proceed via alternative, noncanonical pathways depending on the oncogenic context, with implications for therapy resistance.

Our results show that OIIS has profound consequences for disease progression *in vivo*. OIIS predisposed to tumor relapse from previously regressed tumors, resulting in significantly worse prognosis. These findings extend growing evidence that senescence can paradoxically fuel relapse. Much of our current understanding stems from OIS, initially defined as a potent tumor-suppressive barrier halting premalignant cell proliferation^52,53^. However, accumulating evidence shows that senescent cells can exert pro-tumorigenic effects through SASP-mediated remodeling of the microenvironment and cell-intrinsic adaptations^54–57^. Senescence-associated reprogramming, for instance, promotes stemness and tumor-initiating capacity^21^. In line with this, chemotherapy-induced senescence in acute myeloid leukemia and B-cell lymphoma generates cells that can later re-enter the cell cycle with increased stemness and tumorigenicity^21,22^. Here, we demonstrate that OIIS similarly predisposes tumors to relapse, extending tumor-promoting features of senescence beyond OIS and TIS.

Several mechanisms likely contribute. Previous studies showed that chemotherapy-induced senescence often yields polyploidy, enabling re-entry into the cycle, whereas non-polyploid senescent cells remain arrested^36,58,59^. In our model, relapsed cells displayed extensive chromosomal amplification, a hallmark of advanced, therapy-resistant tumors^60^. Recent work showed that extrachromosomal DNA (ecDNA) generates oncogene dosage heterogeneity affecting adaptation to therapy in MYCN-amplified cancers^61^. Together, these findings highlight how genomic alterations such as polyploidy, chromosomal instability, and ecDNA levels provide routes for tumor cells to persist and adapt under therapeutic pressure. Acquired polyploidy likely facilitates bypass of senescence through activation of alternative oncogenic pathways. Among these, Mdm2 emerged as a key driver. Mdm2 antagonizes p53 and is frequently overexpressed in cancers^62^. Accordingly, pharmacological Mdm2 inhibition with AMG-232 selectively killed relapsed cells, while parental cells remained unaffected. This dependence reflects the fact that, in our model, TagLuc inactivation restores p53 activity, and relapsed cells must therefore re-establish functional p53 suppression to escape OIIS. This aligns with prior work linking p53 reactivation to enforced senescence and growth suppression. For example, in HPV-positive cancers, disruption of the HPV E6–p53 interaction restored p53, induced senescence, and suppressed tumor growth^63^. Similar oncogene-addiction mechanisms have been described in c-MYC-dependent lymphomas, where senescence escape arises through mutations or rearrangements restoring c-MYC activity^41^, while non-degradable c-MYC mutants sustain cell cycle re-entry^24^. In our model, Mdm2 upregulation likely substitutes for TagLuc by suppressing p53 and enabling escape. Interestingly, in other models it could be shown that Mdm2 inhibitors attenuates SASP factors^64^. This dual action, oncogenic suppression and SASP modulation, supports Mdm2 inhibition as a potential therapeutic approach. Nevertheless, our findings are most directly relevant to tumors that retain a functional p53 pathway but acquire Mdm2 overexpression or amplification, a genetic configuration present in several human malignancies. By contrast, in the many human cancers with TP53-mutations, OIIS and its escape are likely governed by p53-independent senescence programs and alternative forms of oncogene addiction, therefore Mdm2 inhibition would not be expected to provide the same therapeutic benefit. Whether analogous oncogene-addiction–like pathways are restored in relapses of TP53-mutant tumors, and whether such alternative senescence-escape circuits can likewise be leveraged therapeutically, will need to be addressed by future studies.

Persistent SASP activation fosters a pro-inflammatory, pro-angiogenic niche that supports regrowth^11,17^. Indeed, analysis of the tumor microenvironment across progression, remission, and relapse revealed marked remodeling of stromal and immune compartments. Relapsed tumors showed increased endothelial cells, suggesting neovascularization, and a shift toward an immunosuppressive milieu characterized by reduced dendritic cells and enriched regulatory macrophages. Comparable remodeling has been reported in TIS models, where SASP factors recruit Gr-1⁺ myeloid cells or polarize macrophages toward M2-like states^54,65,66^. These cells could either blunt immune surveillance or support neoangiogenesis. Our findings show that OIIS similarly reprograms the microenvironment, highlighting that relapse is not solely cell-intrinsic but emerges from co-evolution with the niche. In support of this concept, a recent study combining a next-generation pan-RAS(ON) inhibitor with CDK4/6 inhibitors in experimental PDAC models suggested that therapy-induced senescence can lead to durable tumor control via a senescence-associated immune equilibrium, even though the question, whether tumor-reactive T cells were involved, remained open^26^.

In conclusion, our study reveals that OIIS primes tumors for relapse through polyploidy, chromosomal instability, the acquisition of alternative oncogenic pathways, and SASP-driven microenvironmental remodeling. By characterizing the genetic, metabolic, cytogenetic, and immunological hallmarks of OIIS, we establish it as a distinct form of cellular state that initially constrains tumor growth but ultimately promotes progression, thereby providing a framework for identifying vulnerabilities and risks relevant to preventing cancer relapse.

## METHODS

### Cell culture and oncogene inactivation

Clone 4 cells, TTC #3055 and TetTagLuc cells were cultured in Dulbecco’s modified eagle medium (DMEM, Gibco), supplemented with 10% heat inactivated fetal calf serum (PAN, Biotech) and 50 μg/ml gentamicin (Gibco). Clone 4 and TTC #3055 cells were cultured with 1 μg/ml doxycycline (Sigma) and medium was changed every 3 days. For oncogene inactivation, clone 4 and TTC #3055 cells were cultured, if not otherwise specified, in dox-free medium for 14 days. TetTagLuc cells were cultured in the presence of doxycycline for oncogene inactivation. A375 melanoma cells were cultured in Dulbecco’s modified eagle medium (DMEM, Gibco), supplemented with 10% heat inactivated fetal calf serum (PAN, Biotech) and 1× penicillin-streptomycin (Thermo Fisher Scientific). Cells were treated with 1□μM vemurafenib (TEBU-Bio, in DMSO) for the timepoints indicated in each figure, and medium was changed every 2 days.

### Western blot

Cells were lysed using the mammalian cell lysis kit (Sigma-Aldrich) or the RIPA buffer (Cell Signaling Technology) supplemented with protease/phosphatase inhibitor cocktail (Cell Signaling Technology). Protein concentrations were determined using the BCA assay (Pierce). Proteins were separated by SDS-PAGE gels and transferred to nitrocellulose membranes (Amersham/Bio-Rad). Primary antibodies included anti-SV40 T antigen (2.5 µg/ml, clone PAb416, Calbiochem), p16 INK4A (clones D7C1M and E5F3Y, Cell Signaling Technology), p21 Waf1/Cip1 (clone E2R7A, Cell Signaling Technology). For detection, membranes were incubated with either HRP-conjugated secondary antibodies (Southern Biotech, USA) and visualized using the SuperSignal chemiluminescent substrate kit (Thermo Fisher) on a LUMI-F1 workstation (Roche), or with IRDye 800CW goat anti-mouse IgG and IRDye 680RD goat anti-rabbit IgG (LI-COR Biosciences) and imaged using the LI-COR Odyssey system. Loading controls were performed using anti-β-actin antibodies (0.2 µg/ml polyclonal, Abcam; monoclonal clone AC-15, Sigma-Aldrich).

### SA-**β**-gal staining

Clone 4 cells were cultured in 6-well plates with or without doxycycline until they reached ∼50–80% confluency. A375 cells were seeded at 15,000 cells per well and treated with 1 μM vemurafenib or vehicle (DMSO). Because of their different growth rates, SA-β-gal staining was performed on day 5 for vemurafenib-treated cells and day 2 for DMSO controls. Senescence-associated β-galactosidase activity was detected using either the Senescence Detection Kit (BioVision) or the SA-β-gal Staining Kit (Cell Signaling Technology). In both cases, cells were fixed with the provided fixation solution, washed, and incubated with X-gal staining solution at 37 °C overnight in a dry incubator (no CO₂). After staining, cells were stored in 70% glycerol at 4 °C until imaging. Stained cells were visualized using either an Olympus FSX100 microscope or a DMi8 inverted microscope (Leica Microsystems, 10× objective, scale bar 100 µm) to evaluate SA-β-gal activity.

### Cell cycle analysis

Cells were incubated with 10 μM BrdU (Sigma) for 60 min. Next, cells were washed with PBS and harvested. For permeabilization, cells were resuspended in ice-cold 70% ethanol mixed with PBS and stored for 2 hours at -20°C. Cells were washed again with PBS, incubated with 2 ml 2N HCl/0,5 % Triton X-100 for 30 min, centrifuged and incubated with 2 ml 0.1 M sodium tetraborate (Sigma) for 10 min. After 2 washing steps with PBS, cells were stained with anti-BrdU antibody (Biolegend) supplemented with 0,5% Tween-20 and 1% BSA for 30 min at room temperature. Next, cells were incubated with 20 μg/ml propidium iodide (containing 0,5% Tween-20, 1% BSA and 10 μg/ml RNase A (Qiagen) for 30 min at room temperature. Probes were analyzed via flow cytometry (BD FACS Canto II) and data was analyzed using FlowJo v10.8 Software (BD Life Sciences).

### Proliferation assay

A375 melanoma cells were seeded at 2,000 cells per well into 96-well plates, with separate plates prepared for each time point. On day 0, cells were treated with either vehicle (DMSO) or 1 µM vemurafenib. Proliferation was assessed daily from day 0 to day 4 using the CellTiter 96 AQueous Non-Radioactive Cell Proliferation Assay (MTS; Promega) according to the manufacturer’s instructions. Briefly, the MTS/PMS reagent was added to each well and incubated for 1 h at 37 °C, after which absorbance was measured at 490 nm using a microplate reader. Each condition was plated in triplicate wells, and background absorbance from blank wells was subtracted. For each experiment, the mean absorbance of triplicates per condition was calculated at each time point. The experiment was independently repeated three times on different days, and results were analyzed and graphed using GraphPad Prism v10.

### Real-time quantitative PCR

Total mRNA was extracted from proliferating, TagLuc expressing and senescent, TagLuc non-expressing clone 4 cells using the RNeasy Mini Kit (Qiagen). 1 μg was reverse-transcribed using an oligo-dT primer and ProtoScript II Reverse Transcriptase (NEB) Kit according to the manufacturer’s instructions. The reaction products were diluted (10x) in nuclease free water. 2 μl of the sample was subjected to real-time PCR, which was performed in triplicates using TaqMan probes for the mouse genes CDKN1A, CDKN2A and GAPDH (Applied Biosystems). Expression levels of CDKN1A and CDKN2A relative to that of GAPDH were calculated using the 2^−ΔΔCT^ method. The difference of the cycle threshold (CT) of the target and the reference gene was calculated (ΔCT) and compared between the two samples (ΔΔCT). Finally, expression ratio was calculated according to the formula x = 2^−ΔΔCT^.

### Cytokine antibody array

Proliferating, TagLuc expressing and senescent, TagLuc non-expressing clone 4 cells were cultured in DMEM for 48h to generate conditioned medium (CM). Cells were counted and CM was filtered and analyzed using Mouse Cytokine Antibody Array C2000 (RayBiotech) according to the manufacturer’s protocol. Briefly, array membranes were preincubated with 2 ml of blocking solution. CM was diluted with DMEM to reach volume equivalents of 2 × 10^5^ cells/ml. Array membranes were incubated with 1 ml of CM (3h, room temperature), washed, and incubated with biotin-conjugated antibody cocktail (overnight, 4°C). The next day, HRP-Streptavidin was added (2h, room temperature), followed by a washing step of the array membranes and addition of the detection solution. Chemiluminescence was analyzed using the Lumi Imager F1 (Roche) with an exposure time of 1min.

### ELISA

For the mouse system, conditioned medium (CM) was generated as described above, diluted in DMEM to a volume equivalent of 2 × 10□ cells/ml, and analyzed using IL-6 and MMP-3 ELISA kits (Life Technologies). For the human system, supernatants were directly collected from A375 melanoma cells treated with vemurafenib (1 µM, 15 days) or left untreated, and analyzed using the corresponding human IL-6 and MMP-3 ELISA kits (Life Technologies). All assays were performed according to the manufacturer’s instructions, and absorbance was measured using a µQuant microplate reader (BioTek).

### Luciferase activity assay

Cells were cultured in a 6-well plate, washed with PBS once and incubated with 500 μl of reporter lysis buffer (Promega) for 2 min. A cell scraper was used to harvest the cells and the cell suspension was transferred into Eppendorf tubes. The probes were snap-frozen in liquid nitrogen for 2 minutes, thawed on ice, and centrifuged at 12.000 rpm for 10 min. Protein concentration of the supernatant was determined by NanoDrop (Thermo Scientific). 10 μl of supernatant was mixed with luciferase assay buffer (Promega) and luciferase substrate buffer (Promega), respectively. TagLuc activity was measured using a Mithras LB 940 luminometer (Berthold Technologies) and relative light unit (RLU) was calculated according to total μg protein.

### mRNA sequencing and differential expression analysis

mRNA of the proliferating (TagLuc expressing, +dox) and senescent (TagLuc non-expressing, -dox) cancer cells was isolated using the RNeasy Mini Kit (Qiagen). mRNA integrity was measured using the Agilent Tape Station 4200. Probes were then analyzed via HiSeq4000. Differential gene expression analysis was performed as described before (detailed methods can be found in references^70,71^). In short, sequencing reads were quality-checked and mapped to the mouse genome. Differential expression analysis was performed using the DESeq2 R statistical package. For data analysis, the adjusted p-value was set <0.001 and the minimum log fold change of upregulated genes was set at ≥1. All procedures were performed in triplicates.

### Biological term enrichment analysis

Biological term enrichment analysis was carried out using two different methods. gProfiler2 R package (version 0.2.2) was used to detect enriched pathways (annotated as in the KEGG and Reactome databases) or GO terms among significantly upregulated/downregulated genes^72^. As a complementary approach, gene-set enrichment analysis was carried out using the fgsea R package (version 1.24.0) with cancer state signature genes downloaded from the CancerSEA database^33,73^. For the GSEA analysis, the genes were ranked by decreasing order of the -log10(p-value) multiplied by the sign of the log2-fold-change of each gene. For the results depicted in Extended Figure 4A-B, enrichment of curated gene sets among differentially expressed genes was assessed using a one-tailed hypergeometric test. Up- and down-regulated genes were analysed against the background of all genes in the collection. For each set, the overlap with differentially expressed genes was counted, hypergeometric p-values and odds ratios were computed, and multiple testing correction was applied (FDR, Benjamini–Hochberg). Redundant sets with >90% overlap were pruned, and results were visualized by odds ratio and adjusted p-value, with top sets labeled.

### Oxygen Consumption Rate and Extra-Cellular Acidification Rate Measurements

The oxygen consumption rate (OCR) and extra-cellular acidification rate (ECAR) were measured using a XF96 extracellular flux analyzer (Seahorse Bioscience). Cells were seeded at a density of 8 × 10^4^ (on dox) and 4 × 10^4^ (off dox) cells per well on poly-D-lysine (PDL)-treated plates, each well containing 200 µl of Seahorse XF Base Medium supplemented with 1 mM pyruvate, 2 mM glutamine, and 10 mM glucose. The plates were then incubated in a non-CO_2_ incubator at 37°C for 45 minutes prior to the assay. The optimal cell density to maintain OCR within an optimal range for both conditions was determined experimentally. Basal mitochondrial respiration was assessed by subtracting the non-mitochondrial OCR, measured with 0.5 μM antimycin A and 0.5 μM piericidin A, from baseline OCR. Coupled respiration was determined by subtracting the OCR in the presence of 1.5 μΜ oligomycin A (Sigma) from the basal mitochondrial respiration. Maximal respiration was determined by subtracting the OCR in the presence of 1 μΜ FCCP (Sigma), from baseline OCR. The XF Glycolytic Rate Assay Kit was employed to measure basal and compensatory glycolysis. Basal measurements of extracellular acidification rate (ECAR) were followed by the sequential injection of 0.5 μM rotenone/antimycin A and 50 mM 2-DG. This allowed for the calculation of the glycolytic proton efflux rate (glycoPER), representing both basal glycolysis (in the absence of mitochondrial CO_2_ contribution) and compensatory glycolysis (as a result of electron transport chain inhibition). OCR and glycoPER values were normalized to 2 × 10^4^ cells to facilitate comparison across conditions. Data analysis was performed using the Seahorse Wave desktop software.

### Animal experiments

All animal experiments were performed according to national guidelines and approved by the Landesamt für Gesundheit und Soziales (Berlin, Germany). Mouse strains were housed under specific pathogen-free conditions at the animal facility of the Max Delbrück Center for Molecular Medicine.

For cancer cell transplantation, proliferating or senescent cancer cells were harvested using 0.05% trypsin (Gibco), washed, resuspended in PBS, and mixed with Matrigel (BD, final concentration 10 mg/ml). The suspension was kept on ice until injection. Cells (1 × 10□ or 1 × 10□, as indicated in figure legends) were inoculated subcutaneously (s.c.) into the flanks of mice using a 27-gauge needle (Braun). For oncogene activation, doxycycline (1 mg/ml, Sigma) was administered in light-protected drinking water supplemented with 5% sucrose. To verify TagLuc expression and exclude leakiness in the absence of doxycycline, luminescence imaging was performed *in vitro* prior to transplantation, and *in vivo* bioluminescent imaging (BLI) was carried out as described previously^1^. For the human xenograft experiments, 1 × 10□ A375 melanoma cells that had been pretreated *in vitro* with vemurafenib (1 µM, 14 days) to induce senescence were injected s.c. into Rag2-KO mice. Vemurafenib (TEBU-Bio) was prepared in PBS/PEG300 (7:3), stored at 4 °C, and administered intraperitoneally (25 mg/kg body weight). For the experiments reported here, mice additionally received intraperitoneal injections of the PBS/DMSO (25:1) vehicle as they have been part of a therapy control group.

### Spectral flow cytometry

Subcutaneous tumor plugs were isolated and digested for 1h at 37°C with shaking in a solution of RPMI 1640 medium (Gibco) and collagenase A (1.25 mg/ml). The digested tumors were filtered through a 70µm cell strainer using the plunger of a 5 ml syringe and resuspended in FACS buffer (2% FSC in PBS). After tumor digestion, cells were counted using Countess 3 (Invitrogen). One million cells were transferred into a 96 well U bottom plate, centrifuged 5 min at 350 g and stained with the surface marker panel master mix using FACS buffer and Brilliant Stain buffer (BD) according to manufacturer’s instructions. Cells were stained at 4°C for 30 min, followed by washing with FACS buffer, centrifugation for 5 min at 350 g and resuspension in 100 µL 2% PFA PBS for 15min at room temperature. Cells were washed with FACS buffer, centrifuged for 5 min at 350 g and resuspended in 200 µL FACS buffer for flow cytometric analysis with the Cytek Aurora (Cytek Biosciences). One mouse sample had to be excluded from the analysis due to clogging.

Flow cytometry data was processed as described previously^74^. In brief, raw flow cytometry data was spectrally unmixed using the inbuilt unmixing function of the SpectroFlo (Cytek Biosciences) software. FCS files were imported into FlowJo (BD), parameters for generalized bi-exponential transformation of data were defined for every surface marker individually, and PeacoQC was used as an automatic quality control tool^75^. Live cells were exported using channel values defined by the inbuilt export function of FlowJo, imported into R, and processed with PICtR^74^. In short, data was subsampled per experimental group using the atomic sketching approach implemented in Seurat^76^, clustered using Louvain clustering^77^, and cells were annotated based on their feature expression. Cell type labels were predicted for all cells in the dataset using Linear Discriminant Analysis. Doublets were excluded from the analysis.

### Detection of antigen-specific T cells in the blood

Detection of Tag-specific TCR expression on peripheral blood T cells was achieved by incubation with MHC/peptide tetramers and specific monoclonal antibodies in 50 μl PBS for 15 to 30 min at 4°C. The following antibodies and tetramers were used: monoclonal anti-mouse CD3ε (1:100, clone 145-2C11, Biolegend); monoclonal anti-mouse CD8a (1:100, clone 53−6.7, Biolegend); H-2Kb/ VVYDFLKL (Tag peptide IV) tetramer (1:1000, Biozol/MBL). Cells were washed once with PBS and resolved in 200 μl PBS before analysis using a FACS Canto II or Symphony A1 devices (BD). Erythrocyte lysis was performed with BD lysing solution prior to FACS measurement. BD lysing solution was added for 8 to 10 min followed by one wash with PBS. Data was analyzed with FlowJo software (Treestar, Ashland, Oregon, USA).

### Spectral Karyotyping (SKY)

Spectral Karyotyping was performed as previously described^78^. In brief, metaphase preparation was performed by treating the cells with colcemid for 60 min at a concentration of 0.035 μg/ml. Next, cells were incubated in 75 mM KCl for 20 minutes at 37°C and fixed in a freshly prepared mixture of methanol/acetic acid (3:1) at room temperature. Cell suspension was dropped onto glass slides and SKY images of 8-10 randomly selected metaphase chromosomes per cell line were acquired by staining with a mixture of 5 fluorochromes. Images were captured using an DMRXA epifluorescence microscope (Leica GmbH, Wetzlar, Germany), HCX PL SAPO 63×/1.30 oil objective (Leica), SpectraCube system (Applied Spectral Imaging, Migdal HaEmek, Israel) and analyzed using SKYView imaging software (Applied Spectral Imaging).

### Apoptosis staining

Proliferating (TagLuc expressing, +dox) and senescent (TagLuc non-expressing, - dox) cells were treated with Lys05 at 5, 10 and 20 µM concentration for 7 hours at 37°C. Staining was performed with the apoptosis detection kit (Abcam) according to the manufacturer’s instructions. Briefly, cells were harvested and washed twice with PBS. Next, cells were resuspended in 100 µl binding buffer solution and stained with Annexin V DY-634 and propidium iodide for 15 minutes in the dark. After incubation, 400 µl of binding buffer solution was added and analyzed via flow cytometry (BD FACS Canto II and Symphony A1). Data was analyzed with FlowJo™ v10.8 Software (BD Life Sciences). Additionally, apoptosis was determined on the single cell level by measuring the DNA content of individual cells by flow cytometry. Cellular DNA content was measured with a logarithmic amplification in the FL-2 channel of a FACScan flow cytometer (Becton Dickinson; Heidelberg, Germany) equipped with the CellQuest software. Data are given in percent hypoploidy (i.e., the percentage of cells with a sub-G1 DNA content), which reflects the percentage of apoptotic cells with fragmented genomic DNA. Cells were collected by centrifugation at 2500 rpm, 4°C for 5 min, washed with PBS at 4°C, and fixed in PBS / 2 % (vol/vol) formaldehyde on ice for 30 min, incubated with ethanol/PBS (2: 1, v/v) for 15 min, pelleted and resuspended in PBS containing 40 µg/ml RNase A (Roth; Karlsruhe, Germany). RNA was digested for 30 min at 37°C. Cells were pelleted again and finally resuspended in PBS containing 50 µg/ml PI (Sigma-Aldrich; Taufkirchen, Germany).

### Statistical analysis

All data were analyzed using Prism v10 (GraphPad software) or R v4.4.2. Data are presented as mean ± standard deviation. Statistical analysis of the data was conducted using t-test or one-way analysis of variance (ANOVA). Data comparison with p values of ≤ 0.05 was considered statistically significant. (ns indicates not significant, p > 0.05; * p ≤ 0.05; ** p ≤ 0.01; *** p ≤ 0.001; **** p ≤ 0.0001)

## Data and code availability

The data of the mRNA sequencing, FACS and animal experiments are available upon request from the corresponding author. The raw sequence data reported in the manuscript have been deposited in the Genome Sequence Archive^2^ in National Genomics Data Center^3^, China National Center for Bioinformation / Beijing Institute of Genomics, Chinese Academy of Sciences (GSA: CRA018187) that are publicly accessible at https://ngdc.cncb.ac.cn/gsa. Analysis Code for spectral flow cytometry data can be found at https://github.com/ViktoriaFl/tumor_reject_immune_infiltration/.

## ACKNOWLEDGEMENTS

We thank I. Becker, K. Borgwald and M. Pippow for technical assistance. This work was supported by a grant from the European Union (ERC Advanced Grant Neo-T, number 882963). P.S. is participant in the BIH MD Stipend Program funded by the Charité – Universitätsmedizin Berlin and the Berlin Institute of Health at Charité (BIH). I.M.R. was supported by a Klaus Tschira Boost Fund grant from the German Scholars Organization.

## AUTHOR CONTRIBUTION

P.S., I.M.R. and T.B. conceived the study, analyzed and interpreted the data and wrote the manuscript. P.S., K.H., A.M., K.A., M.T.N., M.V.A, Z.S., L.P, M.T., and I.M.R. performed *in vitro* experiments. P.S., K.H., M.T.N. and M.F.N. performed tumor experiments. V.F., I.S., and S.H. performed and analyzed tumor microenvironment studies using spectral flow cytometry. E.S. generated the SKY images. S.S. performed and analyzed the mRNA sequencing. B.U. performed downstream analysis and visualization of differential expression results and A.A. contributed with compute resources and supervision. G.W. contributed with supervision and technical assistance of in vivo experiments.

---

**Extended Data Figure 1.**
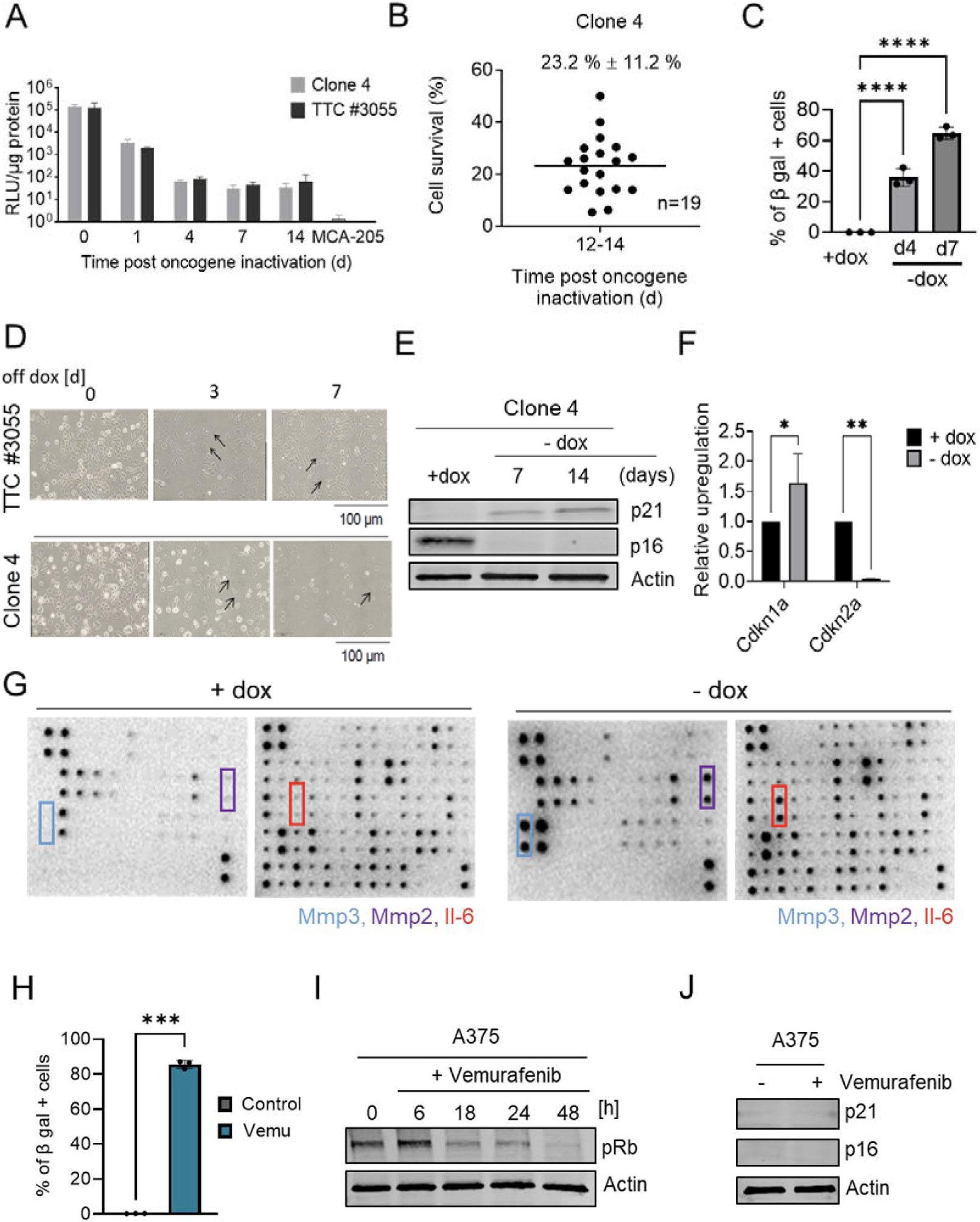
Characterization of senescence and SASP profiles following oncogene inactivation. **A,** Luciferase reporter activity in the presence of dox or at different time points following dox deprivation in clone 4 and TTC #3055 cells. Exposure time: 1s. Error bars indicate standard deviation; n=3 independent experiments. MCA-205 cells serve as negative control. **B,** The viability of clone 4 cancer cells was quantified 12 to 14 days after oncogene inactivation. The percentage of surviving clone 4 cells was calculated relative to the initial cell number before oncogene inactivation; n=19 independent experiments. **C**, Quantification of SA-β-galactosidase-positive cells after four or seven days post-TagLuc inactivation in clone 4 cells. Dara aquired from three independent experiments. Error bars indicate standard deviation. ****p < 0.0001.**D,** Morphological changes of TTC#3055 and clone 4 cells were observed under phase-contrast light microscopy after growth on dox or at three- or seven-days post TagLuc inactivation. Enlarged and flattened cells with irregular shapes are indicated by black arrows. Data from one experiment are shown, although the observed phenotype was consistently observed. **E,** Western blot analysis of p21 and p16 protein levels in clone 4 cells grown on dox or after seven or 14 days post-TagLuc inactivation. β-actin was used as a loading control. Shown is one representative blot out of three independent experiments. **F,** Levels of CDKN1A (p21) and CDKN2A (p16) mRNA relative to GAPDH mRNA in proliferating (TagLuc expressing, +dox) and senescent (TagLuc non-expressing, -dox) clone 4 cells measured by real-time PCR. Error bars indicate standard deviation; n=3 technical replicates. * p < 0,05, ** p < 0,01. **G,** Secretion of cytokines in proliferating (TagLuc expressing, +dox) and senescent (TagLuc non-expressing, -dox) cells. The experiment was conducted once. A comprehensive annotation of the array can be found in Supplementary Document 3. **H**, β-galactosidase staining of A375 cells after Vemu treatment. Percentage of positive cells is shown (mean ± s.d.; n = 3). *** p < 0,001. **I**, Western blot of phosphorylated Rb (pRb) in A375 cells treated with Vemu for the indicated times. Actin served as a loading control. Shown is one representative blot out of three independent experiments. **J,** Western blot showing the expression of the senescence markers p21 and p16 after Vemu treatment. Shown is one representative blot out of three independent experiments.

**Extended Data Figure 2:**
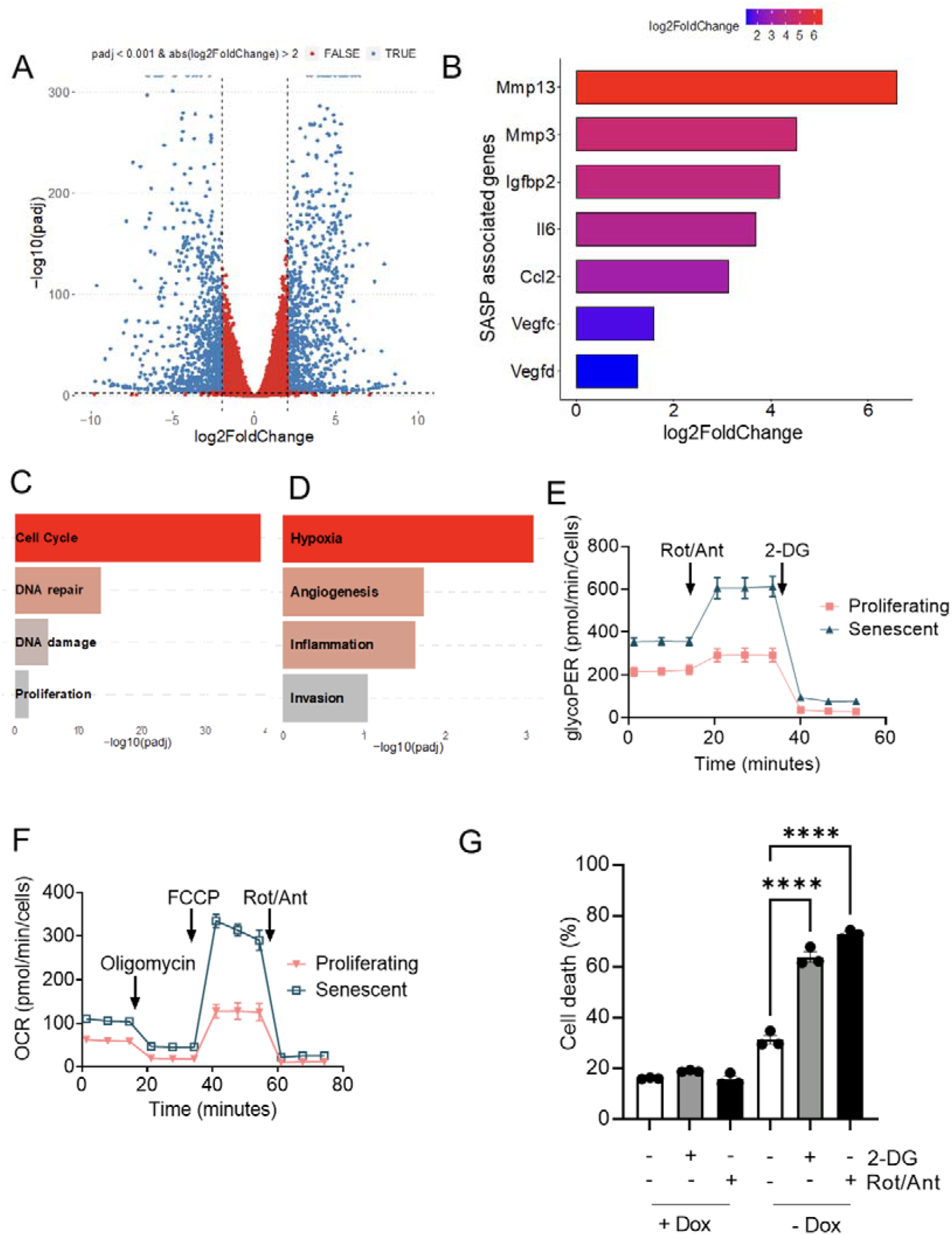
Transcriptional and metabolic changes in senescent *versus* proliferating cancer cells. **A**, Volcano plot showing the differentially upregulated and downregulated genes in senescent (TagLuc-negative, -dox) versus proliferating (TagLuc-expressing, +dox) cells. **B,** Differential expression of SASP-associated genes in senescent (TagLuc-negative, –dox) versus proliferating (TagLuc-expressing, +dox) cells. **C, D,** Gene-set enrichment analysis (GSEA) for CancerSEA cancer-state signatures downregulated (C) and upregulated (D) in senescent (TagLuc-negative, –dox) versus proliferating (TagLuc-expressing, +dox) cells. **E,** Representative profile of the extra-cellular acidification rate after addition of the complex I (rotenone) and complex III (antimycin) inhibitors and the glycolytic inhibitor (2-DG). **F,** Representative profile of the oxygen consumption rate after addition of the complex V inhibitor (oligomycin), the uncoupler (FCCP), and the complex I inhibitor (rotenone) and complex III inhibitor (antimycin) in proliferating (TagLuc-expressing, +dox) and senescent (TagLuc-negative, -dox) cells. **G,** Quantification of cell death in proliferating (+dox) and senescent (–dox, 7 days) cells after treatment with 2-DG (2 mM) or rotenone/antimycin A (R/A, 0.5 µM) for 48 h. Data are shown as mean ± SD (n = 3). \*\*\*\**p* < 0.0001.

**Extended Data Figure 3.**
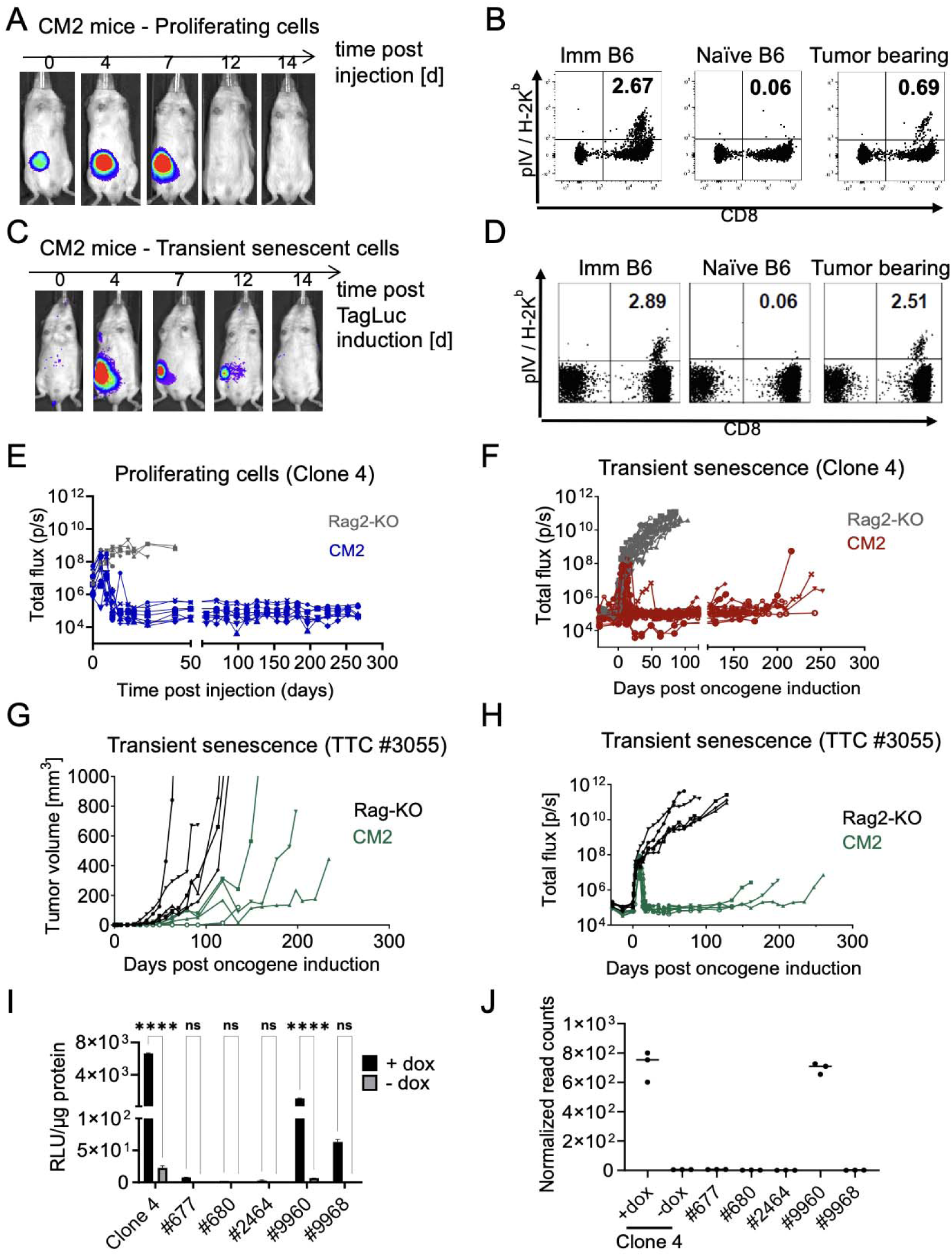
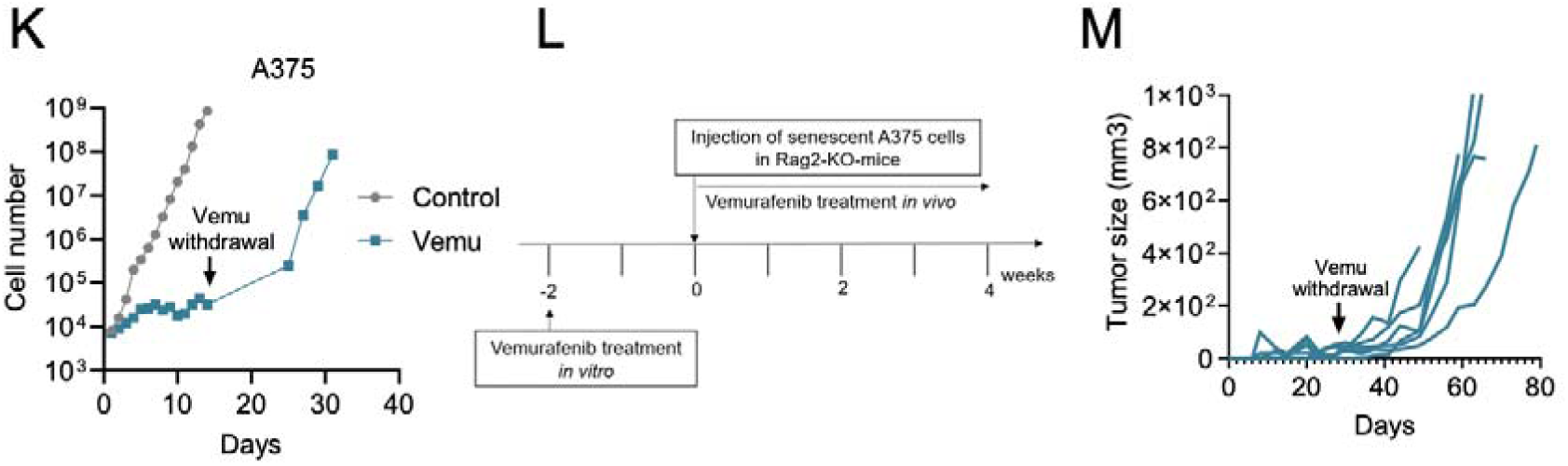
TagLuc-independent growth of relapsed tumors. **A,** Representative CM2 mouse inoculated with proliferating (TagLuc-expressing, +dox) clone 4 cells. TagLuc signal was monitored over time by BL imaging (exposure time: 60 s). **B,** Frequency of Tag-specific CD8⁺ T cells in peripheral blood from Imm B6, naïve B6, or tumor-bearing CM2 mice, measured 10 days after tumor injection. Representative dot plots are shown. **C,** Representative CM2 mouse inoculated with senescent (TagLuc-negative, −dox) clone 4 cells. TagLuc signal was monitored over time by bioluminescence (BL) imaging (exposure time: 60 s) following oncogene reactivation. **D,** Frequency of Tag-specific CD8⁺ T cells in peripheral blood from immunized C57BL/6 (Imm B6), naïve C57BL/6 (Naïve B6), or tumor-bearing CM2 mice, measured 12 days after oncogene re-activation. Representative dot plots are shown. **E,** BL signal kinetics for Rag2-KO (n = 5) and CM2 (n = 8) mice inoculated with proliferating clone 4 cells (TagLuc-expressing, +dox). Data from two independent experiments. **F,** BL signal kinetics for immunodeficient Rag2-KO (n = 10) and immunocompetent CM2 (n = 16) mice inoculated with transient senescent (TagLuc-negative, −dox) clone 4 cells. Data from two independent experiments. **G,** Tumor volume in Rag2-KO (n = 5) mice and relapse growth in CM2 (n = 6) mice following initial rejection of TTC #3055 cells. Data from one experiment. Two CM2 mice died before relapse assessment. **H,** BL signal kinetics for Rag2-KO (n = 5) and CM2 (n = 6) mice inoculated with transient senescent TTC #3055 cells (TagLuc-negative, −dox). Data from one experiment. **I,** Relative light units (RLU) per µg protein for parental clone 4 cells and tumor-derived cell lines from relapsed tumors, cultured ±dox. Four of five relapse-derived cell lines showed no detectable TagLuc activity, even in the presence of dox. Error bars represent standard deviation; n = 2 technical replicates; ns, not significant; **** p < 0.0001. **J,** mRNA sequencing of parental clone 4 and relapse-derived tumor cell lines. Normalized Tag read counts are shown for each sample (n = 3 technical replicates). **K**, A375 cells were treated with 10 µM vemurafenib or left untreated, and cell numbers were monitored over time. Upon drug withdrawal (arrow), cells resumed proliferation. Viable cells were counted by trypan blue exclusion. **L**, Schematic of the *in vivo* experimental setup. Senescent A375 cells were injected into Rag2-KO mice and treated with Lys05 or placebo. **M**, Tumor growth curves of Rag2-KO mice injected with senescent A375 cells and treated with Vemu according to the shown setup.

**Extended Data Figure 4:**
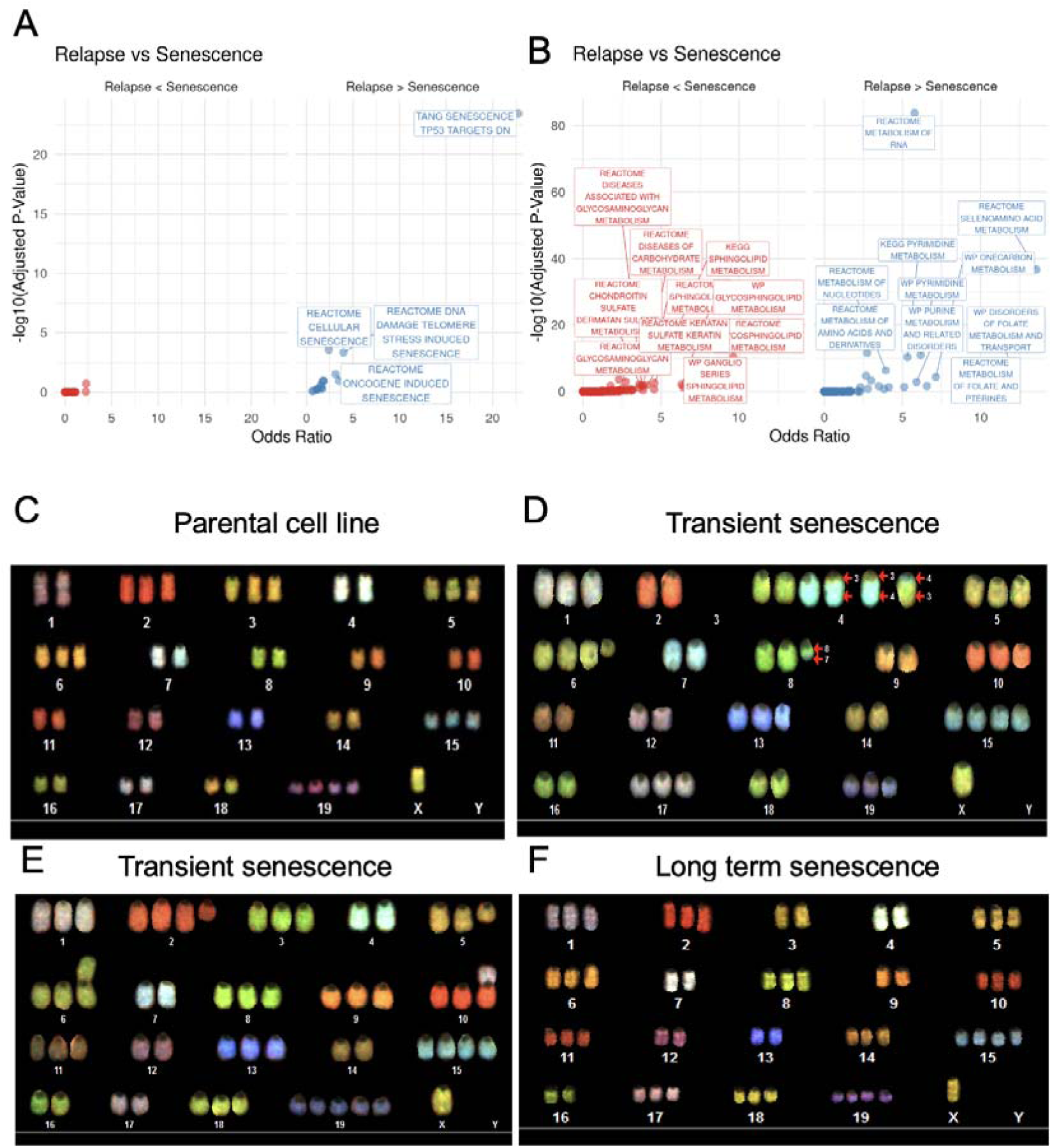
Relapsed tumors are transcriptionally distinct from parental proliferating and senescent cells. **A, B,** Top enriched senescence-related terms (A) and metabolism-related terms (B) detected among genes upregulated (“Relapse > Senescence”) or downregulated (“Relapse < Senescence”) in relapse-derived cancer cell lines compared to both parental proliferating (TagLuc-expressing, +dox) and senescent (TagLuc-negative, – dox) clone 4 cells. Pathway enrichment was based on significantly differentially expressed genes (adjusted p < 0.05, |log₂FC| > 1). **C,** Representative spectral karyotyping (SKY) image of the parental clone 4 cell line maintained under continuous oncogene expression (+dox). **D-F,** Representative SKY images of relapse-derived cell lines from tumors that emerged after transient TagLuc inactivation (D, E) or long-term TagLuc inactivation (F). Images were obtained from 8–10 randomly selected metaphase spreads per cell line, with one representative example shown in each panel.

**Extended Figure 5:**
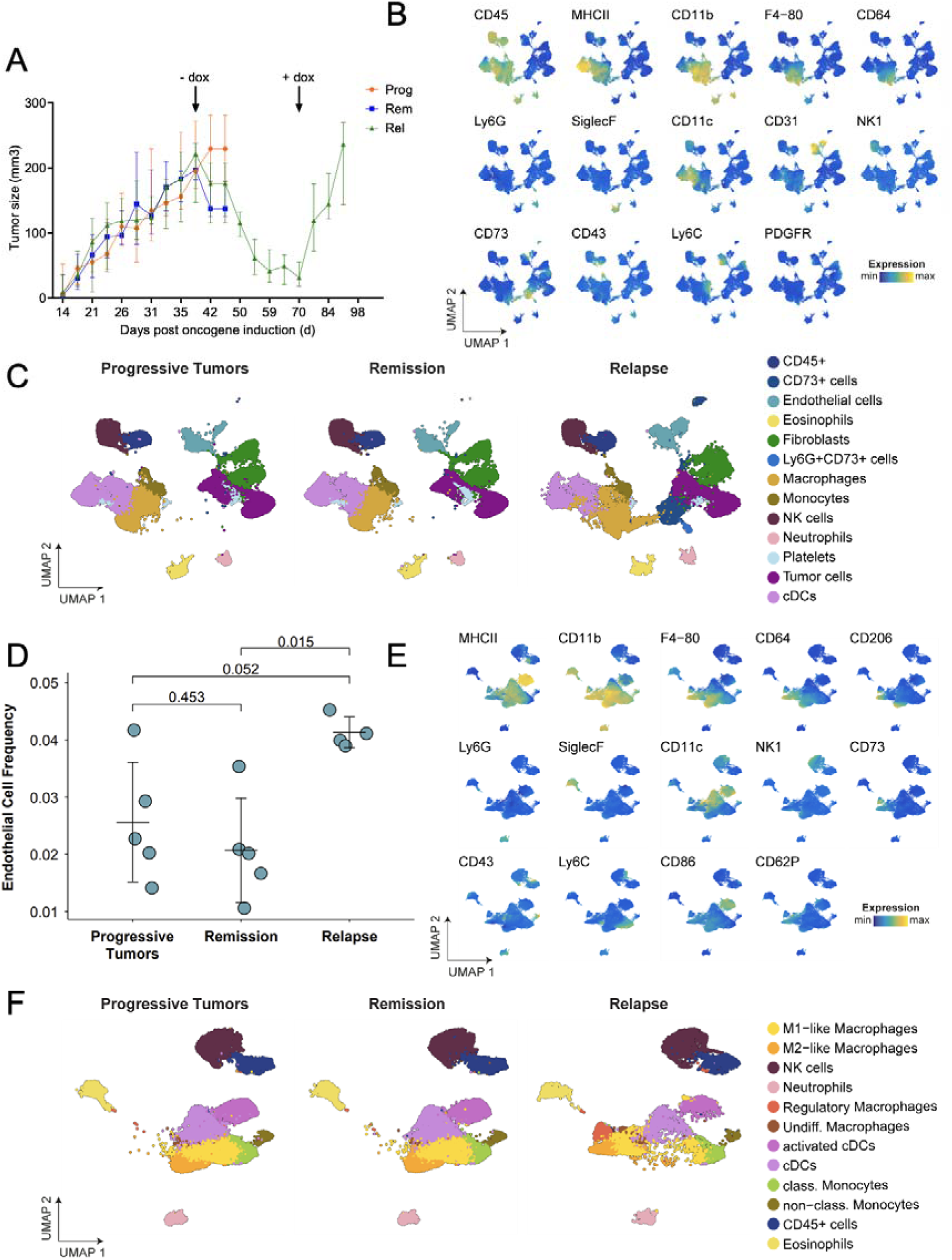
Immune infiltration in progressive tumors, remission and relapse. **A,** Tumor volume in progressive (Prog; n=5), remission (Rem; n=5) and relapse (Rel; n=5) groups depicted in Figure 5B. **B**, UMAP representations from Figure 5D, showing selected feature expression patterns for the overall cellular landscape. **C,** UMAP representation from Figure 5D, split by experimental group. **D,** Quantification of the endothelial cell frequency within all analysed cells across the experimental groups. *P* values were determined with a two-sided Welch’s *t*-test and corrected according to Holm. Error bars indicate the mean and standard deviation. **E,** UMAP representations from Figure 5E, showcasing selected feature expression patterns for the immune cell annotation. **F,** UMAP representation of immune cells (see Figure 5E) split by experimental group. cDCs: conventional dendritic cells, NK cells: natural killer cells, non-class./class. Monocytes: non-classical/classical monocytes, UMAP: Uniform Manifold Approximation and Projection.

